# Yeast-based heterologous production of the Colletochlorin family of fungal secondary metabolites

**DOI:** 10.1101/2023.07.05.547564

**Authors:** Aude Geistodt-Kiener, Jean Chrisologue Totozafy, Géraldine Le Goff, Justine Vergne, Kaori Sakai, Jamal Ouazzani, Grégory Mouille, Muriel Viaud, Richard J. O’Connell, Jean-Félix Dallery

**Affiliations:** Université Paris-Saclay, INRAE, UR BIOGER, 91120 Palaiseau, France; Université Paris-Saclay, INRAE, AgroParisTech, Institut Jean-Pierre Bourgin, 78000 Versailles, France; Centre National de la Recherche Scientifique, Institut de Chimie des Substances Naturelles ICSN, 91190 Gif-sur-Yvette, France

**Keywords:** secondary metabolism, heterologous expression, polycistronic vector, Colletotrichum higginsianum, Saccharomyces cerevisiae.

## Abstract

Transcriptomic studies have revealed that fungal pathogens of plants activate the expression of numerous biosynthetic gene clusters (BGC) exclusively when in presence of a living host plant. The identification and structural elucidation of the corresponding secondary metabolites remain challenging. Here we adapted a polycistronic vector for efficient, seamless and cost-effective cloning of biosynthetic genes using in vivo assembly (also called transformation-assisted recombination) directly in Escherichia coli followed by heterologous expression in Saccharomyces cerevisiae. Two vectors were generated with different auto-inducible yeast promoters and selection markers. The effectiveness of these vectors was validated with fluorescent proteins. As a proof-of-principle, we applied our approach to the Colletochlorin family of molecules. These polyketide secondary metabolites were known from the phytopathogenic fungus Colletotrichum higginsianum but had never been linked to their biosynthetic genes. Considering the requirement for an halogenase, and by applying comparative genomics, we identified a BGC putatively involved in the biosynthesis of Colletochlorins in C. higginsianum. Following the expression of those genes in S. cerevisiae, we could identify the presence of the precursor Orsellinic acid, Colletochlorins and their non-chlorinated counterparts, the Colletorins. In conclusion, the polycistronic vectors described herein were adapted for the host S. cerevisiae and allowed to link the Colletochlorin compound family to their corresponding biosynthetic genes. This system will now enable the production and purification of infection-specific secondary metabolites of fungal phytopathogens. More widely, this system could be applied to any fungal BGC of interest.

## 1. Introduction

Fungi are rich sources of structurally diverse, small molecule natural products, as illustrated by the fact that more than 60% (20,304) of the 33,372 microbial natural products currently catalogued in the Natural Products Atlas were isolated from fungi (https://www.npatlas.org, May 2023) (van Santen et al., 2022). Well-known for their uses in medicine or agriculture, fungal natural products play important roles in the adaptation of fungi to their ecological niches, for example as toxins to compete with other microorganisms, as protection against environmental stresses, or in the case of pathogens, as effectors to facilitate the infection of plant or animal hosts (Collemare et al., 2019; Oberlie et al., 2018).

Fungal genes involved in the biosynthesis of natural products are typically located side-by-side in the genome as so-called biosynthetic genes clusters (BGCs). These clusters usually contain genes encoding one or two key enzymes that generate the backbone of the molecule and a varying number of genes encoding accessory enzymes that decorate the initial molecule. Genes coding for transporters or transcription factors can also form part of the BGC (Keller, 2019). This colocalization in the genome facilitates the identification of BGCs, and several bioinformatic tools (e.g. SMURF, antiSmash, MIBiG, CusProSe) have been developed to mine the ever-increasing number of sequenced fungal genomes released in public databases (Blin et al., 2021; Khaldi et al., 2010; Oliveira et al., 2023; Terlouw et al., 2023). Analysis with these tools has shown that a single fungal genome can contain more than 80 non-redundant BGCs (Han et al., 2016; Inglis et al., 2013; Liang et al., 2018; Valero-Jiménez et al., 2020). However, for the vast majority of these predicted BGCs, the chemical products are currently unknown.

Colletotrichum higginsianum is a plant-pathogenic ascomycete fungus that causes disease on various cultivated Brassicaceae as well as the model plant Arabidopsis thaliana (O’Connell et al., 2004). Resequencing the genome of C. higginsianum revealed the presence of 77 non-redundant BGCs, of which only 14 (18%) can be linked to known chemical products based on their similarity to BGCs characterized in other fungi (Dallery et al., 2017; O’Connell et al., 2012). Transcriptional analysis showed that 19 C. higginsianum BGCs are induced at particular stages of plant infection. Among these, 14 are specifically expressed during the initial biotrophic phase of penetration and growth inside living plant cells, two are upregulated during later necrotrophic growth in dead tissues, while three are expressed at both infection stages (Dallery et al., 2017; O’Connell et al., 2012). These clusters are poorly expressed, or not at all, by axenic cultures of C. higginsianum, which has hindered isolation of the corresponding fungal metabolites in sufficient amounts for determination of their chemical structures and analysis of their biological activities.

Various strategies can be used to activate the expression of cryptic fungal BGCs in axenic cultures. For example, variation of the culture conditions (media composition, static/liquid cultures), as in the ‘One strain many compounds’ (OSMAC) technique (Bode et al., 2002; Hewage et al., 2014) and co-cultivation with other microbes (Yu et al., 2021). Other approaches include over-expressing global or cluster-specific transcriptional activators (Chiang et al., 2009; von Bargen et al., 2013), and opening up chromatin structure by the chemical or genetic manipulation of epigenetic regulator proteins, such as CCLA, KMT6 and LAEA (Lyu et al., 2020; Pimentel-Elardo et al., 2015). In C. higginsianum, we previously deleted the CCLA subunit of the COMPASS protein complex, which mediates mono-, di- and trimethylation of lysine 4 in histone H3. Cultures of the resulting mutant over-produced a number of terpenoid compounds belonging to the Higginsianin, Colletorin/Colletochlorin and Sclerosporide families (Dallery et al., 2019).

Although these genome-wide strategies have been successfully used to isolate new compounds from some fungi, they are untargeted and do not activate all silent BGCs. Heterologous expression provides a way to activate specific BGCs of interest and facilitates the large-scale production of metabolites in axenic cultures. In this approach, an entire BGC is cloned into a heterologous microbial host that can be easily cultivated (Ahmed et al., 2020; Bond and Tang, 2019; Chiang et al., 2013; Gomez-Escribano and Bibb, 2011; Pfeifer et al., 2001; Zhang et al., 2018). For heterologous production of fungal secondary metabolites, the most frequently used hosts have been Escherichia coli (Kealey et al., 1998; Pfeifer Blaine et al., 2003), the yeasts Saccharomyces cerevisiae and Pichia pastoris (Cochrane et al., 2016; Gao et al., 2013; Kealey et al., 1998; Xue et al., 2017) and some filamentous fungi, including species of Aspergillus, Trichoderma and Penicillium (Heneghan et al., 2010; Nielsen et al., 2013; Pohl et al., 2020; Shenouda et al., 2022). S. cerevisiae has the advantage of being easily genetically manipulated, grows rapidly in liquid culture and can support the protein folding and post-translational modifications occurring in filamentous fungi. Yeast also produces very few endogenous secondary metabolites, which facilitates the purification and isolation of heterologous compounds (Bond et al., 2016; Ishiuchi et al., 2012; Yu et al., 2013; Zhao et al., 2020).

To synchronously activate all the genes in a biosynthetic pathway during heterologous expression it is often necessary to laboriously change the native promoter and terminator of each gene (Harvey et al., 2018; Pahirulzaman et al., 2012). To avoid this, Hoefgen et al. (2018) designed an expression vector that allows the concerted expression of multiple genes as a single polycistron, where all the genes are placed under the control of a single promoter, allowing their co-expression. The self-splicing porcine teschovirus P2A DNA sequence is inserted at the 3’ end of each gene of the polycistron, which induces bond skipping by the ribosome during translation, thereby releasing the individual proteins (Kim et al., 2011). The vector also incorporates a split fluorescent reporter gene so that correct translation of the polycistronic transcript can be monitored by microscopy. The vector was successfully used to transfer the psilocybin biosynthetic pathway into A. nidulans (Hoefgen et al., 2018).

In the present study, we have adapted the polycistronic vector of Hoefgen et al. (2018) for the heterologous expression of fungal BGCs in baker’s yeast. To validate this expression system, we chose the Colletochlorins and their non-chlorinated counterparts, the Colletorins, a well-characterized family of prenylated polyketide compounds produced by C. higginsianum (Dallery et al., 2019). Using genome mining and comparative genomics, we identified a candidate BGC that could be involved in this family of molecules. Four genes, encoding a polyketide synthase, a non-ribosomal peptide synthase-like enzyme, a putative prenyltransferase and an halogenase, were expressed in S. cerevisiae. The resulting yeast culture supernatants were found to contain the expected products and intermediates of this biosynthetic pathway, namely Orsellinic acid, Colletorins B and D, Colletochlorin B and D and Colletorin D acid.

## 2. Materials and Methods

### 2.1. Biological material and growth conditions

A summary of the Saccharomyces cerevisiae strains used in this study is given in Supplementary File 1. In vivo assembly and propagation of plasmids was performed using the E. coli TOP10 strain. Bacteria were maintained and propagated in LB broth, supplemented with antibiotics and agar as required. The yeast strains were cultivated at 28°C in the selective medium YNB (cat. no. Y0626, Sigma), supplemented with drop-out without leucine (cat. no. Y1376, Sigma), without uracil (cat. no. Y1501, Sigma) or without both leucine and uracil (cat. no. Y1771, Sigma), and supplemented with 2% glucose and 2% agar. For the heterologous production of metabolites, the yeast strains were cultivated in YPD liquid medium (yeast extract 10 g·L, peptone 20 g·L and glucose 20 g·L) for 72 h at 30°C with agitation on a rotary shaker at 250 rpm.

### 2.2. Yeast expression vector construction

All vectors developed in the present study were built by in vivo assembly (IVA) of DNA fragments in E. coli. The PCR primers used to amplify the fragments were designed to provide a minimum of 22 bp of overlapping sequence between adjacent fragments for homologous recombination in the bacteria. The amount of fragment used for IVA varied according to the fragment size: 200-300 bp: 1.5 pmol, 300-600 bp: 1.0 pmol, 600-1000 bp: 0.5 pmol, 1000-3000 bp: 0.2 pmol, 3000-5000 bp: 0.1 pmol, > 5000 bp: 0.05 pmol. Residual plasmid matrix in the PCR reactions was removed by digestion at 37°C for 2 h with 10U of DpnI enzyme before proceeding further. All the cloning fragments were obtained by PCR amplification using Q5 polymerase according to the manufacturer’s instructions (cat. no. M0491L, New England Biolabs). Diagnostic PCR was performed using GoTaq polymerase (cat. no. M7805, Promega).

The plasmid pHYX104 was constructed by amplifying by PCR the polycistronic fragment (VenusN-P2A-TEV-P2A-VenusC) from pV2A-T (Hoefgen et al., 2018) using primers P953 and P954, the URA3 marker gene, 2µ origin, AmpR and ColE1 origin from pNAB-OGG (Schumacher, 2012) with primers P949 and P950, the ScADH2 promoter with primers P951 and P952 and the ScHIS5 terminator with primers P955 and P956. The four fragments were assembled by IVA in E. coli. The resulting plasmid pHYX104 was amplified by PCR with primers P963 and P964 to remove the AmpR gene and replace it with the modified KanR gene amplified with primers P965 and P966 from pV2A-T to take advantage of the SwaI/PmeI restriction sites. The resulting plasmid pHYX105 was further modified with an EcoRV-free version of the URA3 gene obtained by gene synthesis to give rise to pHYX138.

The plasmid pHYX106 was constructed by amplifying the polycistronic fragment (VenusN-P2A-TEV-P2A-VenusC) from pV2A-T with primers P953 and P954, the LEU2 marker gene from M4755 (Voth et al., 2003) with primers P971 and P972, the 2µ origin, KanR and ColE1 origin from pHYX105 with primers P967 and P950, the ScPCK1 promoter with primers P975 and P976 and the ScPRM9 terminator with primers P977 and P978. The five fragments were assembled by IVA in E. coli. The resulting pHYX106 was further modified by PCR with primers P1204 and P1205 to insert a single nucleotide mutation to remove the EcoRV restriction site present in the LEU2 marker gene, giving rise to the plasmid pHYX137 (Supplementary File 2).

The plasmid pHYX163 was constructed by digesting the plasmid pHYX153 with SwaI and the pHYX154 with PmeI following the strategy described by Hoefgen et al. (2018). The digestion was performed overnight (nearly 16h). After heat inactivation, the digested fragments (0.1 pmol of each fragment of interest) were assembled by IVA in E. coli. The resulting plasmid pHYX163 was digested by PmeI and assembled with the plasmid pHYX152 previously digested with SwaI, to obtain the pHYX164 plasmid. The plasmid pHYX172 was derived from plasmid pCfB2312, and was constructed by amplifying pCfB2312 with primers P1312 and P1313, and the prenyltransferase (codon-adapted for yeast) from pHYX154 with primers P1310 and P1311. The two fragments were gel-purified and assembled by IVA in E. coli. All primers used are listed in Supplementary File 3. The following plasmids were deposited with Addgene: pHYX137 (#202814), pHYX138 (#202815), pHYX143 (#202816) and pHYX173 (#202817).

### 2.3. Modifications of the yeast chassis strain

The yeast strain S. cerevisiae BJ5464-NpgA was modified by inserting the Botrytis cinerea NADPH cytochrome P450 reductase gene BcCPR1 (Bcin12g03180) at the yeast locus XI-3 as described by Mikkelsen et al. (2012). Competent yeast cells were prepared according to Knop et al. (1999). The CRISPR-Cas9 transformation was performed as described by Jessop-Fabre et al. (2016) using the plasmids pCfB2312, pCfB3045 and XI-3-bccpr1 yielding the strain BJNBC (Supplementary File 1).

### 2.4. Comparative analyses of the biosynthetic gene cluster 16

The protein sequences of the genes belonging to the BGC16 of C. higginsianum IMI 349063 (CH63R_05468 to CH63R_05483) as predicted in Dallery et al. (2017) were used to query the NCBI nr database using cblaster v1.3.9 (Gilchrist et al., 2021) with the `search` module and default parameters but limiting the investigations to fungi with `-eq “txid4751[orgn]”` for the Colletochlorin part of BGC16, or limiting the investigations to Colletotrichum spp. with the parameter `-eq “txid5455[orgn]”` for the entire BGC16. The BGCs identified were retrieved with the module `extract_clusters` and used as input for clinker software v0.0.23 using default parameters (Gilchrist and Chooi, 2021). Manual editing of the cartoons was performed to represent the contig ends where appropriate.

### 2.5. Cloning of reporter genes and biosynthetic genes

The gene predictions of CH63R_05468 to CH63R_05471 were manually inspected for correctness using RNA-Seq datasets previously published (O’Connell et al., 2012). The CDS of CH63R_05468 appeared to be composed of six exons instead of only five in the initial prediction. Each gene was then synthesized without stop codons and with optimization of the codons for S. cerevisiae and exclusion of common restriction enzyme sites using the GenSmart™ Codon Optimization tool (Genscript Biotech B.V., Netherlands). The optimized sequences can be found in Supplementary File 4. Each coding sequence was individually cloned into the EcoRV linearized pHYX137 and subsequently assembled together following the strategy described by Hoefgen et al. (2018) using SwaI and PmeI restriction enzymes followed by IVA in E. coli. The gene coding mScarlet-I was amplified from the plasmid Double UP mNeongreen to mScarlet (Addgene #125134) by PCR using primers P1214 and P1215 and cloned into the EcoRV linearized pHYX137 by IVA in E. coli. The gene coding Tobacco Etch Virus (TEV) protease was synthesized (Genscript Biotech B.V., Netherlands), then amplified by PCR using primers P1177 and P1178 and cloned into the EcoRV linearized pHYX137 by IVA in E. coli. All coding sequences were verified by Sanger sequencing after their initial cloning and their presence was further verified by PCR after subsequent combination of vectors for multigene expression. A summary of the plasmids constructed in this study is presented in Supplementary File 5.

### 2.6. Yeast transformation

Established protocols were used for the transformation of plasmids into yeast strain BJNBC and the preparation of frozen competent yeast cells (Knop et al., 1999). To obtain the strain BJNBC015, yeast transformation was performed in two steps. First, the plasmids pHYX164 and pHYX172 were integrated, yielding the strain BJNBC014, and then the plasmid pHYX173 was integrated into strain BJNBC014, giving the strain BJNBC015.

### 2.7. Microscopy

For confocal microscope observations, yeast strains were cultivated in YPD media at 28°C for 2 days. For staining nuclei, samples were fixed with 4% formaldehyde in PBS for 30 min, spun down and rinsed once in PBS, permeabilized with 0.2% Triton X-100 in PBS for 5 min and rinsed three times in PBS. Samples were then incubated in 15 µg·mL of DAPI (4’,6-diamidino-2-phenylindole) in PBS for 30 min, washed for 5 min and mounted on microscope slides prior to observation. Samples were imaged by sequential scanning using a Leica TCS SPE laser scanning microscope (Leica Microsystems) equipped with an APO 40× (1.15 NA) oil immersion objective. Venus, mScarlet-I and DAPI were excited using the 488 nm, 532 nm and 405 nm laser lines, respectively.

### 2.8. Protein extraction

For total protein extraction, yeast strains were cultivated in the appropriate YNB medium until OD reached 0.4. The cultures were then centrifuged (3,000 × g) for 10 min and the pellets resuspended in fresh YNB medium supplemented with 2% (w/v) glucose and 3% (v/v) ethanol. Yeast cells were pelleted by centrifugation at 3,000 × g for 5 min at 4°C and immediately resuspended in lysis buffer (8 M urea, 5% [w/v] SDS, 40 mM Tris-HCl pH 6.8, 0.1 mM EDTA, 0.4 mg·mL bromophenol blue) supplemented with 1% (v/v) β-mercaptoethanol, 1× protease inhibitor cocktail (cat.no. P8215, Sigma-Aldrich), 5 µg.mL leupeptin (cat.no. L2884, Sigma-Aldrich) and 1 mM PMSF (cat. no. P7626, Sigma-Aldrich). PMSF was renewed every 7 min until the samples were frozen at -80°C or loaded on a gel. Immediately after resuspension, glass beads were added up to the meniscus and the mixture incubated at 70°C for 10 min. Cells were disrupted using a vortex mixer for 1 min and debris were pelleted by centrifugation at 18,000 × g for 5 min at 4°C. After the supernatants were recovered on ice, 75 µL of lysis buffer was added to the pellets, boiled at 100°C for 5 min, centrifuged again and the supernatants were finally combined with those from the first centrifugation.

### 2.9. Immunoblot assay

Proteins were separated by SDS-PAGE on 4-15% gradient Mini-Protean TGX Stain-free gels (cat. no. 4568083, Bio-Rad), transferred onto PVDF membranes (cat. no. 1704273, Bio-Rad) and subsequently blocked with 5% (w/v) BSA in TBST buffer. The membranes were incubated for 1 h at RT with a mouse anti-2A primary antibody (cat. no. MABS2005, Merck) diluted 1:2000 in 1% (w/v) BSA in TBST. The membranes were then rinsed 15 min in TBST then 3 x 5 min in TBST before incubation for 1 h at ambient temperature with HRP-coupled goat anti-mouse secondary antibody diluted 1:5000 (cat.no. ab6728, Abcam). The membranes were then rinsed with TBST as above before chemiluminescence detection using the Clarity Western ECL substrate kit (cat. no. 1705060, Bio-Rad). Gels and blots were recorded with a ChemiDoc Imaging System (Bio-Rad).

### 2.10. General chemistry procedures

The cultivation of yeast strains harboring each plasmid and isolation of the chemical compounds were as described previously by Harvey et al. (2018). After preculturing the yeast strains in selective medium supplemented with 2% (w/v) glucose (2 days at 28°C with shaking at 250 rpm), the preculture (10 mL) was inoculated into 1 L of YPD medium in a 2 L Erlenmeyer flask (total 4 × 1L) and incubated for 72 h at 30°C with shaking at 250 rpm. The YPD culture was then centrifuged aseptically (5000 × g, 5 min) and the supernatant was incubated overnight with sterile XAD-16N resin (Dow Chemicals) for solid phase extraction (Dallery et al., 2019). The resin was collected by filtration and extracted for 2 h in ethyl acetate (100 mL) followed by 2 h in methanol (100 mL). Lyophilized cell pellets were resuspended in acetone (3 × 30 mL), sonicated for 3 × 15 min, centrifuged at 5000 × g, 5 min between each acetone addition, followed by extraction with methanol (3 × 30 mL). Ethyl acetate extracts were dried over anhydrous sodium sulphate. Similar extracts were pooled, evaporated under reduced pressure and resuspended in HPLC grade methanol. The crude extracts were then analyzed on an Alliance 2695 HPLC instrument equipped with a 2998 photodiode array, a 2420 evaporative light scattering and an Acquity QDa mass detector (Waters Corporation). The HPLC column used was a 3.5 µm C-18 column (Sunfire 150 × 4.6 mm) operating a linear gradient from H_2_O to CH_3_CN, both containing 0.1% formic acid, for 50 min at 0.7 mL·min . Thin layer chromatography plates (Si gel 60 F 254) were purchased from Merck. Purified standards of Orsellinic acid, Colletorin D, Colletorin D acid, Colletochlorin D, Colletochlorin B and Colletochlorin A were dissolved at 1 mg·mL^-1^ in methanol. All standards were purified as previously described (Dallery et al., 2019) except Orsellinic acid that was purchased from ThermoFisher Scientific (cat.no. 453290010).

### 2.11. Untargeted analysis of different colletochlorin derivatives

Untargeted analysis was performed using a UHPLC system (Ultimate 3000 Thermo) coupled to quadrupole time of flight mass spectrometer (Q-Tof Impact II Bruker Daltonics).

Separation was performed on an EC 100/2 Nucleoshell Phenyl-Hexyl column (2C×C100Cmm, 2.7Cμm; Macherey-Nagel) at 40°C, with a flow rate of 0.4 mL·min, for 5 μL injected. The mobile phases used for the chromatographic separation were: (A) 0.1% formic acid in H_2_O; and (B) 0.1% formic acid in acetonitrile. Elution was as follows: 5% phase B for 2 min, the gradient elution increased linearly to 50% phase B in 13 min, followed by a further linear increase to 100% phase B in 10 min, then 100% phase B for 3 min and the final gradient linear elution decreased to 5% phase B for 7 min.

Data-dependent acquisition methods were used for mass spectrometer data in negative ESI mode using the following parameters: capillary voltage, 4.5CkV; nebulizer gas flow, 2.1Cbar; dry gas flow, 6CL·min ; drying gas in the heated electrospray source temperature, 200°C. Samples were analysed at 8CHz with a mass range of 100–1500Cm/z. Stepping acquisition parameters were created to improve the fragmentation profile with a collision RF from 200 to 700CVpp, a transfer time from 20 to 70Cμsec, and collision energy from 20 to 40CeV. Each cycle included a MS fullscan and 5 MS/MS CID on the 5 main ions of the previous MS spectrum.

### 2.12. Data processing of untargeted metabolomic data

The .d data files (Bruker Daltonics) were converted to .mzXML format using the MSConvert software (ProteoWizard package 3.0; Chambers et al., 2012). mzXML data processing, mass detection, chromatogram building, deconvolution, samples alignment and data export were performed using MZmine-2.37 software (http://mzmine.github.io/) for negative data files. The ADAP chromatogram builder (Myers et al., 2017) method was used with a minimum group size of scan 9, a group intensity threshold of 1000, a minimum highest intensity of 1000 and m/z tolerance of 10Cppm. Deconvolution was performed with the ADAP wavelets algorithm using the following setting: S/N threshold 10, peak duration rangeC=C0.01–2Cmin RT wavelet range 0.02–0.2Cmin, MS2 scan were paired using a m/z tolerance range of 0.05CDa and RT tolerance of 0.5Cmin. Then, isotopic peak grouper algorithm was used with a m/z tolerance of 10Cppm and RT tolerance of 0.1min. All the peaks were filtered using feature list row filter keeping only peaks with MS2 scan. The alignment of samples was performed using the join aligner with an m/z tolerance of 10Cppm, a weight for m/z and RT at 1, a retention time tolerance of 0.2Cmin. Metabolites accumulation was normalized according to the weight of dried extract for the relative quantification. Molecular networks were generated with MetGem software (Olivon et al., 2018; https://metgem.github.io) using the .mgf and .csv files obtained with MZmine2 analysis. The molecular network of ESI− datasets was generated using cosine score thresholds of 0.60.

### 2.13. Metabolite annotation of untargeted metabolomic data

Metabolite annotation was performed in three consecutive steps. First, the obtained RT and m/z data of each feature were compared with our library containing the 6 standards based on their RT and m/z. Second, the ESI− metabolomic data used for molecular network analyses were searched against the available MS spectral libraries (Massbank NA, GNPS Public Spectral Library, NIST14 Tandem, NIH Natural Product and MS-Dial), with absolute m/z tolerance of 0.02, 4 minimum matched peaks and minimal cosine score of 0.60. Third, not-annotated metabolites that belong to molecular network clusters containing annotated metabolites from steps 1 and 2 were assigned to the same chemical family and annotation was carried out on the basis of MS/MS spectrum comparisons.

## 3. Results

### 3.1. Heterologous expression vectors and modification of the yeast recipient strain

In their study, Hoefgen et al. (2018) described the polycistronic plasmid pV2A-T designed to express multiple secondary metabolism (SM) genes under the control of a single promoter. In this polycistronic system each gene is separated by a TEV-P2A sequence. The P2A sequence encodes a self-cleaving peptide releasing the upstream protein with a 33 amino acids tail in C-term and the downstream protein with a proline in N-term. The TEV peptide is recognized and cut by the TEV-protease enzyme, reducing the C-term tail to 6 amino acids. Each polycistron contained the VenusN and VenusC genes on the first and last position of the coding sequence, respectively. Because both genes contain a nuclear localization signal (NLS), when the polycistronic transcript is translated, the VenusN and VenusC proteins accumulate in the yeast nucleus, where they self-assemble to produce a yellow fluorescent protein. The presence of this fluorescence in the nucleus is thus an indicator of the production of the polycistronic proteins. By digestion with the EcoRV restriction enzyme, biosynthetic genes from the cluster of interest can be introduced between the VenusN and VenusC genes. Polycistronic plasmids containing the desired genes can then be fused after digestion by SwaI or PmeI (Hoefgen et al., 2018).

Here, we generated the plasmids pHYX137 and pHYX138, both of which can be used in S. cerevisiae (Figure 1; Supplementary File 2). Both vectors harbour auto-inducible promoters from yeast, namely pPCK1 or pADH2, respectively, which are repressed in the presence of glucose and activated after the diauxic shift during ethanol-anaerobic fermentation (Harvey et al., 2018). This allows to disconnect biomass accumulation from SM production, which is a valuable feature when the SM are toxic. Both promoters are poorly induced in selective medium, in contrast to rich medium (Lee and DaSilva, 2005).

**Figure 1:**
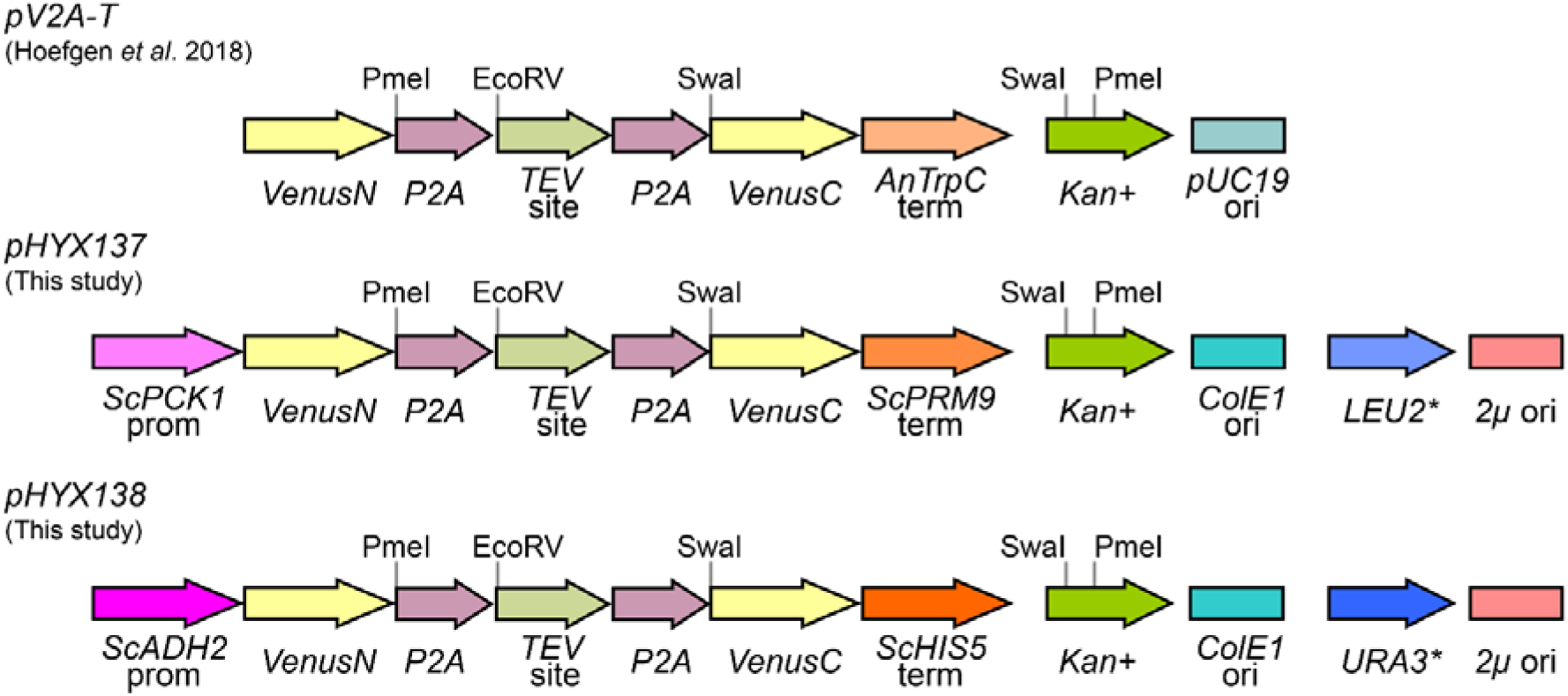
Features of the pV2A-T plasmid described by Hoefgen et al. (2018) and of the two plasmids, pHYX137 and pHYX138, adapted for S. cerevisiae expression, described in this study. Complete maps are shown in Supplementary File 2.

In addition to the polycistronic gene, the plasmids pHYX137 and pHYX138 possess a yeast 2µ origin of replication and the pUC19 origin of replication was also replaced by the ColE1 origin. The nutritional selection genes, either LEU2 or URA3, were included and silent mutations were introduced to remove the EcoRV restriction sites. This allows to linearize the pHYX137 or pHYX138 with EcoRV and to clone individually the coding sequences by in vivo assembly (IVA) in E. coli, which is a fast, cost-effective and simple method. Likewise, the coding sequences are successively assembled in a single polycistron by digesting the plasmids with either SwaI or PmeI and direct transformation of E. coli with the unpurified plasmid fragments for IVA.

To enhance the production of heterologous polyketides in the yeast S. cerevisiae, the strain BJ5464-npga was previously modified by the deletion of two vacuolar proteases, PEP4 and PRB1 (Lee et al., 2009) and the introduction of the npgA gene (Bond et al., 2016) involved in activating the ACP domain of PKS enzymes. Here, we used the strain BJNBC, a derivative of BJ5464-npga that we generated in the frame of a project involving BGCs with numerous cytochrome P450 enzymes. The modification involved the integration of an NADPH-cytochrome reductase from a filamentous fungus. This NADPH-cytochrome reductase was successfully used to enhance the heterologous production of B. cinerea abscisic acid in S. cerevisiae (Otto et al., 2019).

### 3.2. Fluorescence-based validation of the yeast heterologous expression system

To verify the proper transcription of the polycistronic gene, and correct translation and separation of the individual proteins, a gene coding for the mScarlet-I red fluorescent protein was introduced between the coding sequences of the VenusN and VenusC genes. The mScarlet-I gene was first introduced into plasmid pHYX137, giving the plasmid pHYX143, which was then transformed into the BJNBC yeast strain. The transformed yeast was cultivated 24 h in YNB medium supplemented with 2% (w/v) glucose without leucine. Epi-fluorescence microscopy revealed that the yellow fluorescence of Venus was present in the yeast nucleus, where it colocalized with the blue fluorescent DNA stain DAPI, whereas the red fluorescence of mScarlet-I, which lacked an NLS, was distributed through both the cytoplasm and nucleus (Figure 2). These observations confirm that proteins encoded by the polycistronic gene had been well-transcribed and separately translated in yeast, and that Venus was correctly assembled in the yeast nucleus from the two complementary non-fluorescent fragments.

**Figure 2:**
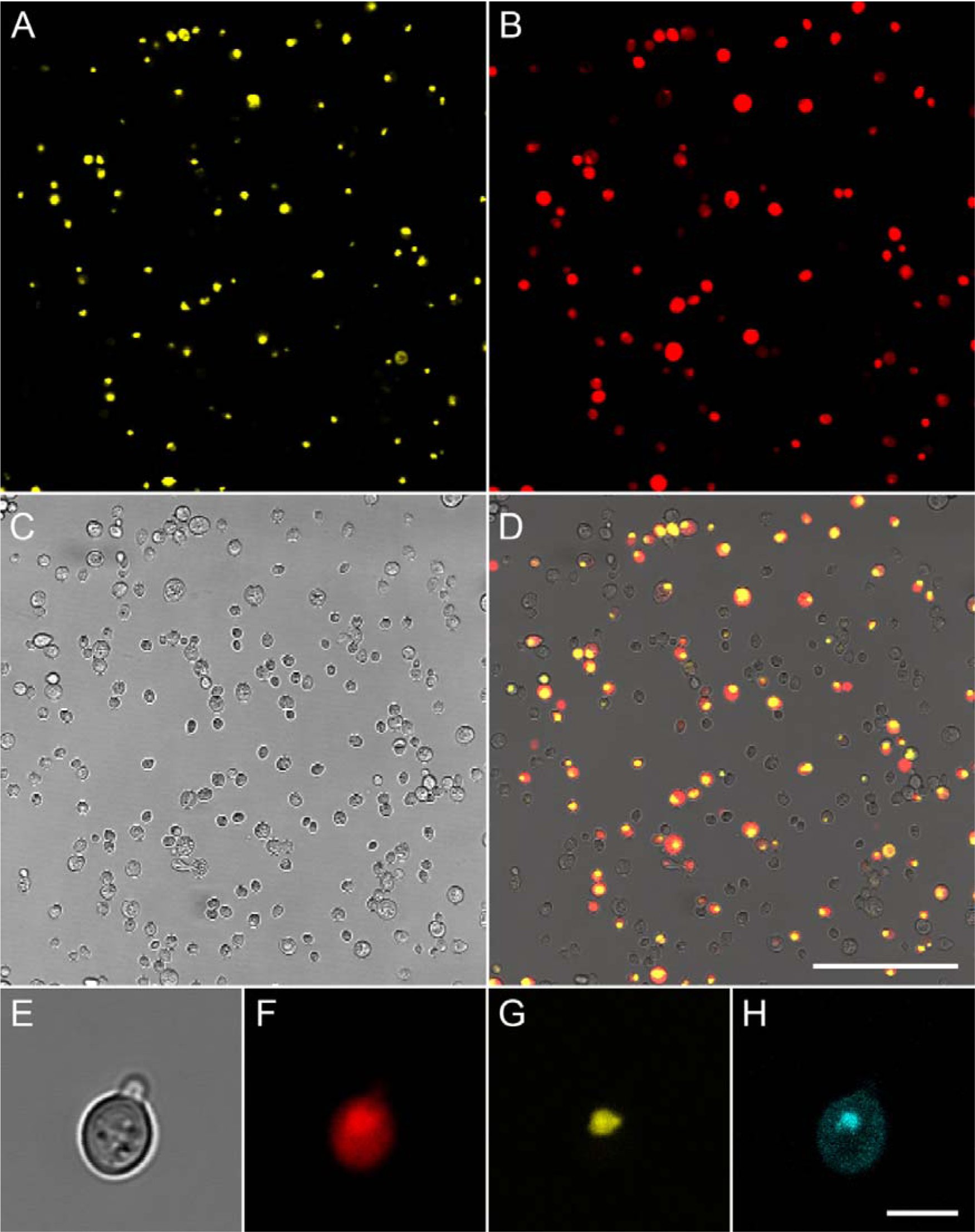
Confocal microscopy of the yeast strain BJNBC-003 expressing Venus-NLS and mScarlet-I fluorescent proteins and stained with DAPI to detect DNA. A-D, Bar= 50 µm. E-H, from left to right, bright-field image showing the yeast cells, red fluorescence corresponding to mScarlet-I, yellow fluorescence corresponding to Venus and, blue fluorescence corresponding to DAPI. The mScarlet-I signal is distributed throughout the cell whereas the Venus signal is co-localized with DNA in the nucleus. Bar = 5 µm.

### 3.3. Selection of a biosynthetic pathway to test the expression system

Previously, we isolated several members of the Colletochlorin family of secondary metabolites from C. higginsianum and we proposed an hypothetical biosynthetic pathway based on the isolated molecules and the plausible expected enzymatic activities (Dallery et al., 2019). In order to identify the biosynthetic gene cluster (BGC) responsible for producing Colletochlorins, we looked for putative halogenases (InterPro signature IPR006905) in the C. higginsianum IMI 349063 genome. The only BGC with an IPR006905 signature was BGC16, comprising genes CH63R_05468 to CH63R_05483 in the original prediction (Dallery et al., 2017). Recently, Tsukada et al. (2020) reported the heterologous production of Higginsianins as well as other decalin-containing diterpenoid pyrones by expressing 8 of the 16 genes in BGC16. None of the Higginsianins are chlorinated and only one PKS (ChPKS11) was required for the biosynthesis of Higginsianins despite the presence of a second PKS (ChPKS10) in the cluster, suggesting the BGC16 comprises two BGCs side-by-side or intertwined. To test this hypothesis, we examined the conservation of BGC16 in 57 genome-sequenced Colletotrichum spp. (NCBI taxid 5455) using the cblaster tool (Figure 3). Interestingly, the BGC16 was found in six species belonging to four different species complexes, namely C. higginsianum and C. tanaceti (Destructivum complex), C. musicola and C. sojae (Orchidearum complex), C. spaethianum (Spaethianum complex) and C. chlorophyti. In C. tanaceti and C. sojae, the BGC16 homologous genes were found respectively on two and three different contigs with each part being located at contig ends, suggesting problems of genome assembly rather than locations on different chromosomes. Interestingly, in C. chlorophyti only the genes required for making Higginsianin-like molecules were retrieved. Homologues of the genes CH63R_05468 to CH63R_05472 were absent from the C. chlorophyti NTL11 genome and were hypothesized to be involved in the biosynthesis of Colletochlorins (Figure 3).

**Figure 3:**
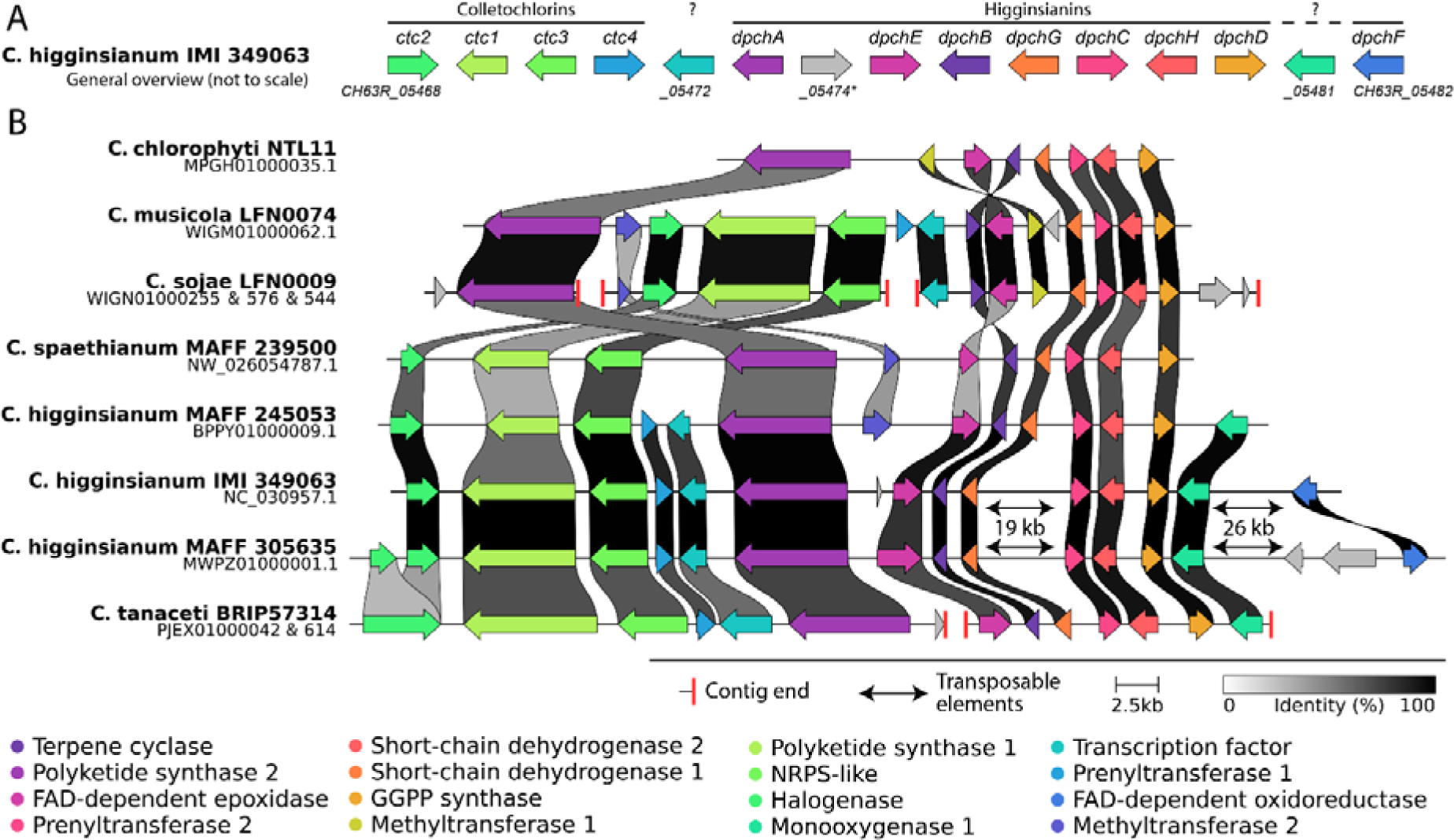
General overview of the biosynthetic gene cluster BGC16 (A) and its conservation and microsynteny in genome-sequenced Colletotrichum spp. (B). The BGC is actually composed of two BGCs side-by-side or intertwined, one for biosynthesis of Colletochlorins and the other for biosynthesis of Higginsianins. The Higginsianin genes (dpch) were characterized by Tsukada et al. (2020). The gene CH63R_05474 is a pseudogene whereas the gene CH63R_05472 encodes a predicted transcription factor. When present, regions composed of repeated transposable elements are shown as double-headed arrows together with their length. Vertical red bars denote contig ends. Fungal BGCs are often misassembled during genome sequencing and split over several contigs due to difficult-to-assemble long stretches of repeats. The intensity of grey/black shading represents the percentage of amino acid identity.

Using the cblaster tool, we investigated the conservation of genes putatively responsible for Colletochlorins biosynthesis (CH63R_05468 to CH63R_05472) in other fungi. Clustered gene homologues were found mostly in Sordariomycetes belonging to the Glomerellaceae, Nectriaceae, Hypocreaceae and Stachybotryaceae families (Figure 4). Only three Eurotiomycetes had homologous BGCs (Aspergillus ellipticus and two Talaromyces spp.). None of the four homologous clusters found in Stachybotrys species contained an halogenase-encoding gene. Consistently, Stachybotrys bisbyi cultures produced only non-chlorinated prenylated derivatives of Orsellinic acid, notably LL-Z1272β, also called Ilicicolin B (Li et al., 2016). Among the retrieved homologues, we also found the BGC in Acremonium egyptiacum responsible for biosynthesis of Ascochlorin, another prenylated yet chlorinated derivative of Orsellinic acid (Figure 4). Based on these findings and knowledge of the experimentally-determined biosynthetic route for LL-Z1272β and Ascochlorin, as well as the predicted pathway for Colletochlorins, we selected genes CH63R_05468 (ctc2, halogenase), CH63R_05469 (ctc1, also known as ChPKS10), CH63R_05470 (ctc3, also known as ChNRPS-like04) and CH63R_05471 (ctc4, prenyltransferase) for heterologous expression in S. cerevisiae.

**Figure 4:**
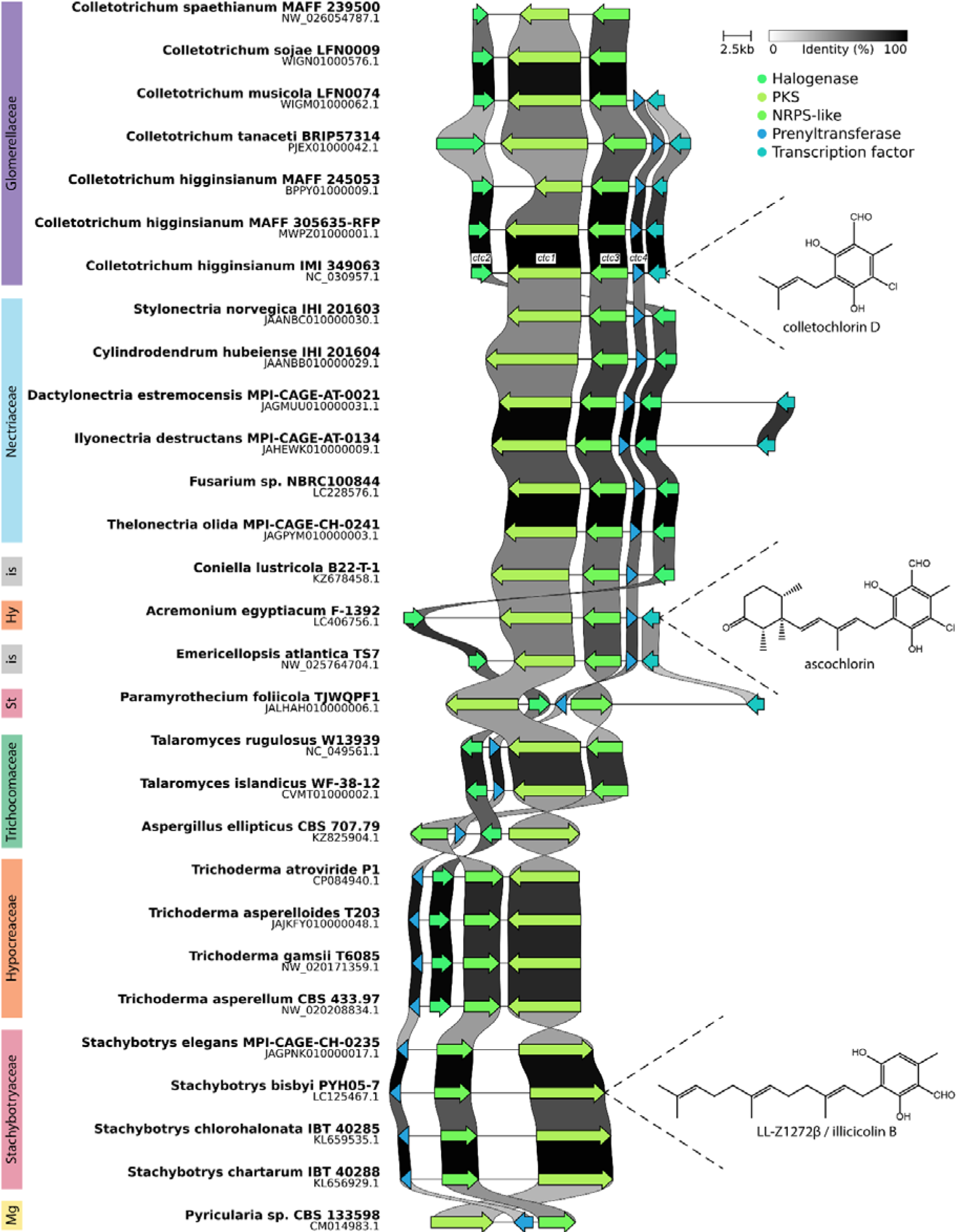
Conservation and microsynteny of the Colletochlorin (ctc) biosynthetic genes cluster in fungi. In Colletotrichum higginsianum IMI 349063, this BGC was initially predicted to be a single BGC16 with the dpch BGC responsible for Higginsianins biosynthesis, located side-by-side. Genes are color-coded according to their putative or experimentally-confirmed functions. The intensity of grey/black shading represents the percentage of amino acid identity. Experimentally-verified BGCs are shown with one of their molecular products. Hy, Hypocreaceae; is, incertæ sedis; Mg, Magnaporthaceae; St, Stachybotryaceae.

### 3.4. Heterologous production of Orsellinic acid in yeast

After careful examination of the gene models and intron borders using available RNA-Seq data, the coding sequences (CDS) of ctc1 to ctc4 were codon-optimized for S. cerevisiae and de novo-synthesized. Each CDS was then cloned individually into the plasmid pHYX137. A polycistron containing the four ctc CDS under control of the PCK1 promoter of S. cerevisiae was constructed by successive digestions with either SwaI or PmeI followed by in vivo assembly (Transformation-Assisted Recombination, TAR) directly in E. coli. This plasmid was named pHYX164 and was transformed into the adapted yeast strain BJNBC, giving BJNBC-008. All plasmids and strains used in this study are described in Supplementary File 1 and 5.

The yeast strains BJNBC-001 (containing the empty polycistronic plasmid pHYX137) and BJNBC-008 (harboring the Colletochlorin gene cluster in pHYX164) were cultured for three days and then metabolites were extracted and analysed by HPLC. Only one molecule was detected in BJNBC-008 that was not present in BJNBC-001. This molecule had a retention time of 16.5 min, a molecular weight of 168 and a UV spectrum with maxima at 227, 259 and 298 (Figure 5). These same characteristics were also shown by an orsellinic acid (OA) standard, indicating that the molecule detected in culture extracts is OA (Figure 5). The presence of OA, the first molecule in the proposed biosynthetic pathway, but none of the other expected molecules, suggests that the polyketide synthase was functional but not the prenyltransferase (Figure 6).

**Figure 5:**
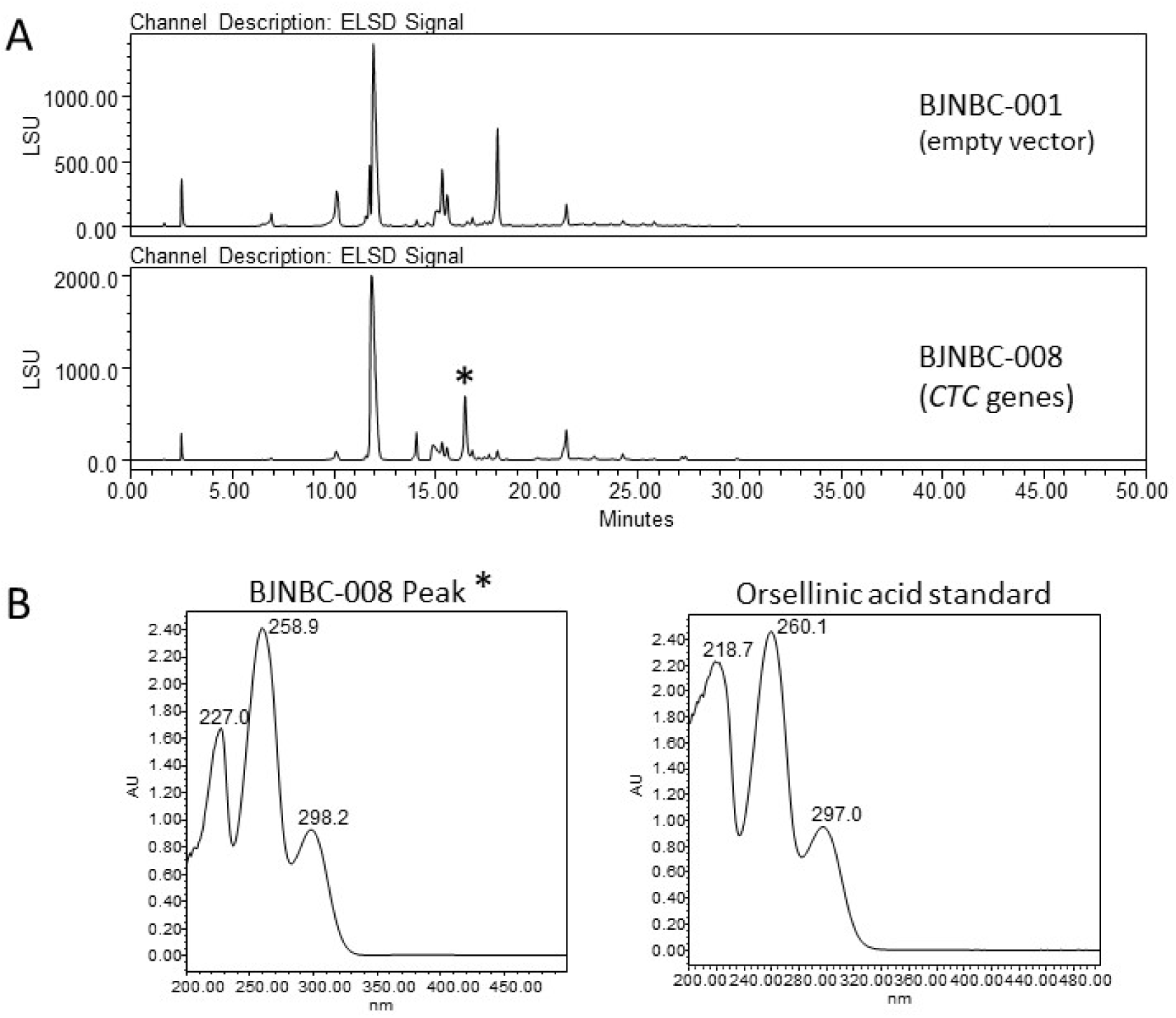
Monitoring the production of Colletochlorin biosynthetic intermediates using HPLC-PDA-ELSD-MS. (A) ELSD chromatogram of crude extracts of strains BJNBC-001 (empty vector) and BJNBC-008 (ctc1 to ctc4 genes expressed from a polycistron). Only Orsellinic acid could be detected among the known intermediates of the Colletochlorin family. (B) UV spectra of the differential peak identified in BJNBC-008 (asterisk) and of the Orsellinic acid standard.

**Figure 6:**
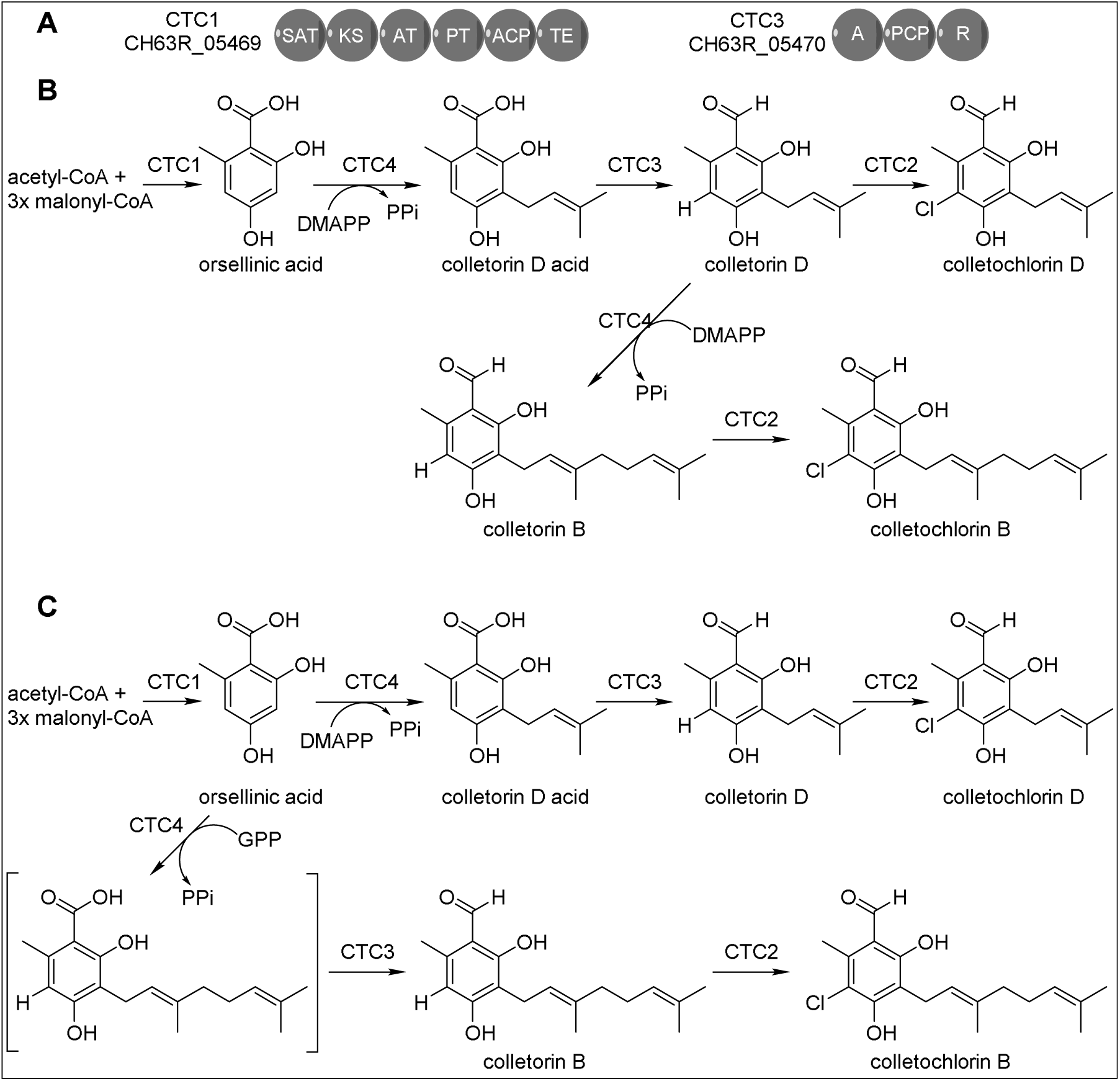
Proposed biosynthetic pathways of the Colletochlorins. (A) Domain structure of CTC1 and CTC3 proteins. (B) Hypothetical scenario 1 where the CTC4 prenyltransferase accepts only DMAPP (dimethylallylpyrophosphate) as isoprene donor. (C) Hypothetical scenario 2 where the CTC4 prenyltransferase accepts both DMAPP and GPP (geranylpyrophosphate) as isoprene donors. Domains: ACP, acyl-carrier protein; AT, acyl transferase; KS, ketosynthase; PT, product template; SAT, starter-unit acyltransferase; TE, thioesterase; A, adenylation; PCP, peptidyl-carrier protein; R, reduction.

All the enzymes expressed from this polycistron retain a P2A tag at their carboxyl terminus. To verify that all the enzymes in the pathway were present in the transgenic yeast BJNBC-008, we used immunoblotting with antibodies raised against the P2A peptide (Hoefgen et al., 2018). Protein samples were collected at two time-points, the first (t0) corresponded to when the culture reached an OD_600_ of 0.4 and the culture medium was changed to a new one containing 2% glucose and 3% ethanol. At that point, the polycistronic gene under control of the PCK1 promoter was expected to be repressed, as PCK1 is repressed in glucose-containing media. The second time-point (t24) corresponds to 24 h after t0. At t24, all the glucose was supposed to be consumed by the yeast and ethanol fermentation had started (Lee and DaSilva, 2005), thus activating the polycistronic gene expression.

Before induction (t0), a band at 29kDa corresponding to the VenusN protein was detected in both BJNBC-001 and BJNBC-008. The detection of VenusN in non-induced conditions (t0) showed that the polycistron was expressed at a very low level in these conditions. After induction (t24h), only the VenusN protein was found in BJNBC-001, whereas in the BJNBC-008 yeast, four bands were detected at 29, 61, 119 and 233 kDa, corresponding to VenusN, CTC2, CTC3 and CTC1, respectively (Figure 7). However, the prenyltransferase CTC4 (expected Mr = 40 kDa) was not detectable. This apparent absence of the prenyltransferase could explain why only the first molecule in the pathway, Orsellinic acid, was obtained from BJNBC-008 cultures.

**Figure 7:**
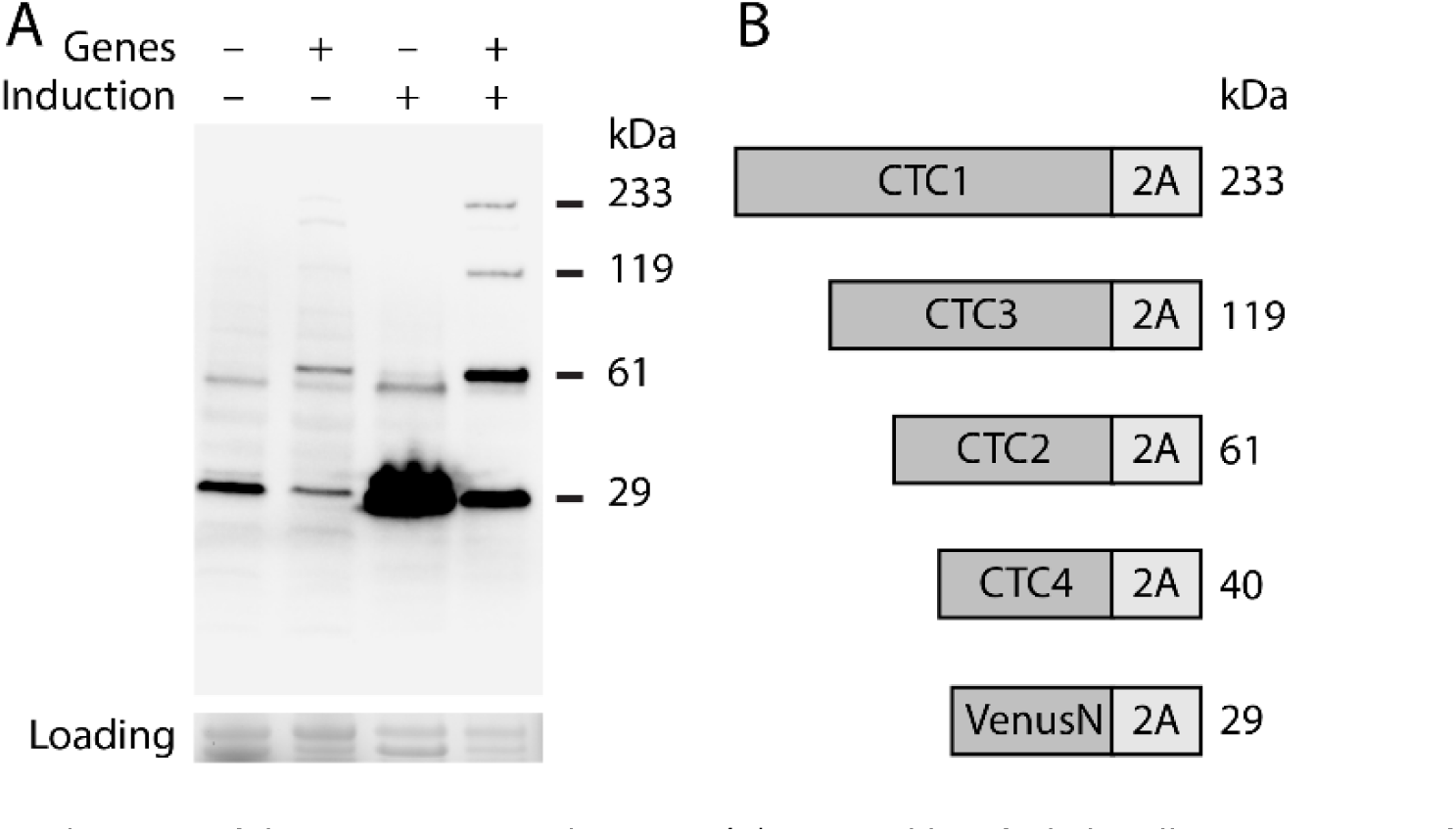
Immunodetection of the CTC proteins and VenusN. (A) Immunoblot of whole-cell protein extracts from the strains BJNBC001 (empty vector) and BJNBC008 (ctc genes) at t0 (optical density of 0.4; repressive medium replaced by inductive medium) and t24 after induction. The proteins were detected with an anti-2A antibody. Note that VenusN is present also in the empty vector. Equal loading was assessed using TGX Stain-Free gels. (B) Schematic representation of the expected proteins with their size.

### 3.5. Heterologous production of the Colletochlorin metabolites in yeast

To overcome the absence of the CTC4 protein in the BJNBC-008 protein extract, a new plasmid pHYX172 was made containing only the prenyltransferase gene under the control of the strong and constitutive TEF1 promoter. Another plasmid (pHYX173) containing the gene encoding the TEV protease was also introduced into the yeast strain BJNBC, which allows cleavage of the C-terminal P2A-tail from the polycistronic enzymes. Two new yeast strains were generated: BJNBC-015 containing all three plasmids pHYX164 (polycistron with Colletochlorin genes), pHYX173 (polycistron with TEV protease gene) and pHYX172 (ctc4 alone), and as a control, the strain BJNBC-017 containing the pHYX137 and pHYX138 empty vectors. Metabolites were extracted from 3-day-old cultures and then analysed by LC-QToF-MS.

The expected molecular ions corresponding to Colletorins and Colletochlorins were readily found in samples from BJNBC-015 and with retention times and masses similar to those for the purified standards Orsellinic acid (RT, 6.40; m/z 167.0344 [M-H]), Colletorin D acid (RT, 13.51; m/z 235.0970 [M-H]), Colletorin D (RT, 15.22; m/z 219.1021 [M-H]), Colletochlorin B (RT, 20.02; m/z 321.1257 [M-H]) and Colletochlorin D (RT, 17.06; m/z 235.0631 [M-H]) (Figure 8).

**Figure 8:**
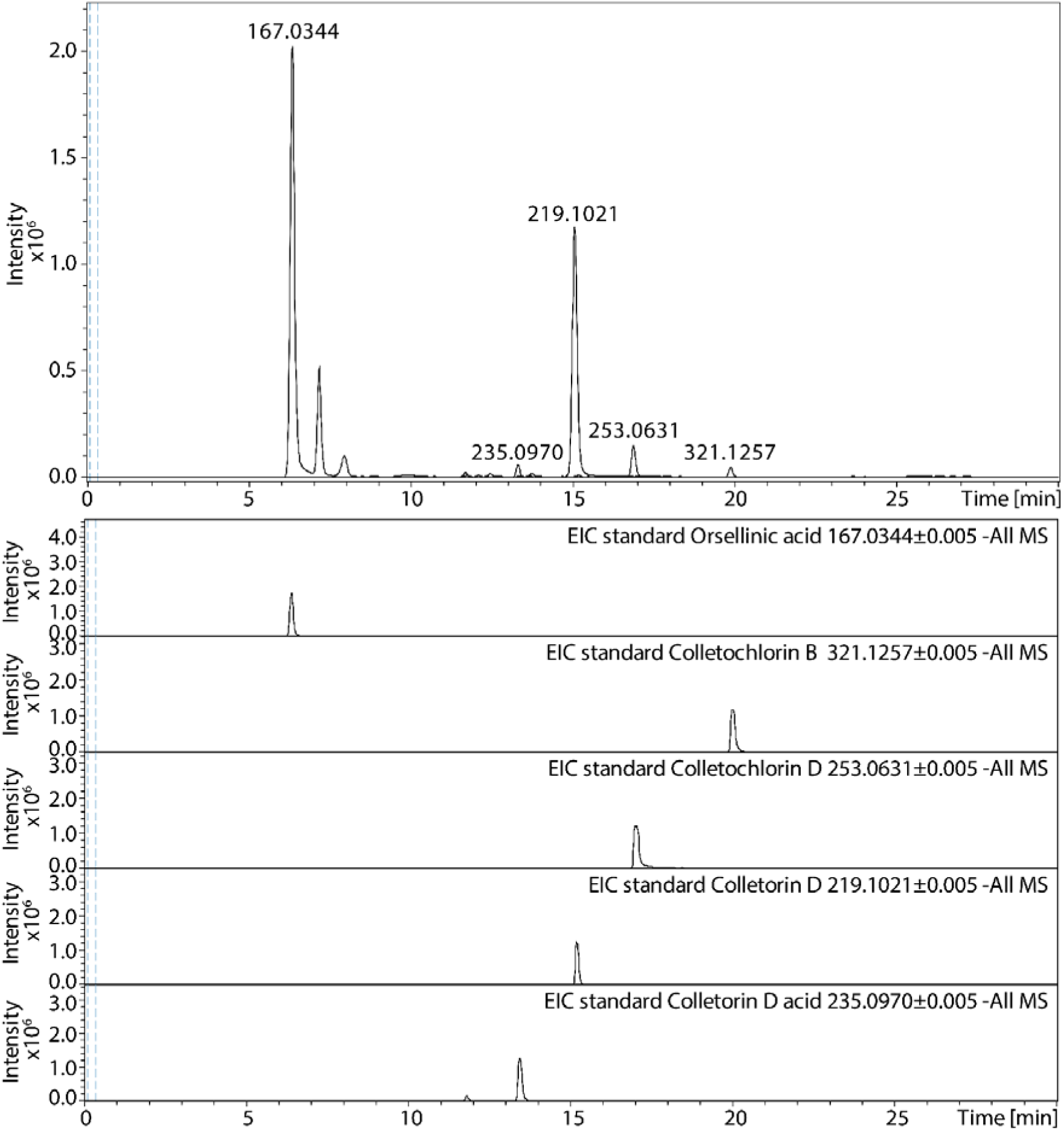
Combined extracted ion chromatograms (EIC) of the culture supernatant of the strain BJNBC015 expressing all four ctc genes in a polycistron and an additional copy of ctc4 (prenyltransferase) with its own promoter. EIC of standards of the different biosynthetic intermediates in the Colletochlorin pathway are also represented.

Next, we performed a non-targeted analysis, comparing the BJNBC015 extract with the control BJNBC-017 extract. This confirmed the detection of all the standard molecules previously described in the BJNBC-015 extract. In addition, another molecule (RT, 18.71; m/z 287.1666 [M-H]) was detected. The molecular mass of this molecule corresponds to that of Colletorin B, the non-chlorinated form of Colletochlorin B. In order to validate this hypothesis, we carried out a comparative analysis of the fragmentation pattern of Colletochlorin B and Colletorin B (Figure 9; Supplementary File 6). Taken together, the results show that Orsellinic acid, Colletorin D acid, Colletorin B and D, and Colletochlorin B and D were detected in the yeast harbouring genes ctc1 to ctc4 of C. higginsianum BGC16, and validated that the proposed gene cluster does indeed encode the Colletochlorin biosynthetic pathway.

**Figure 9:**
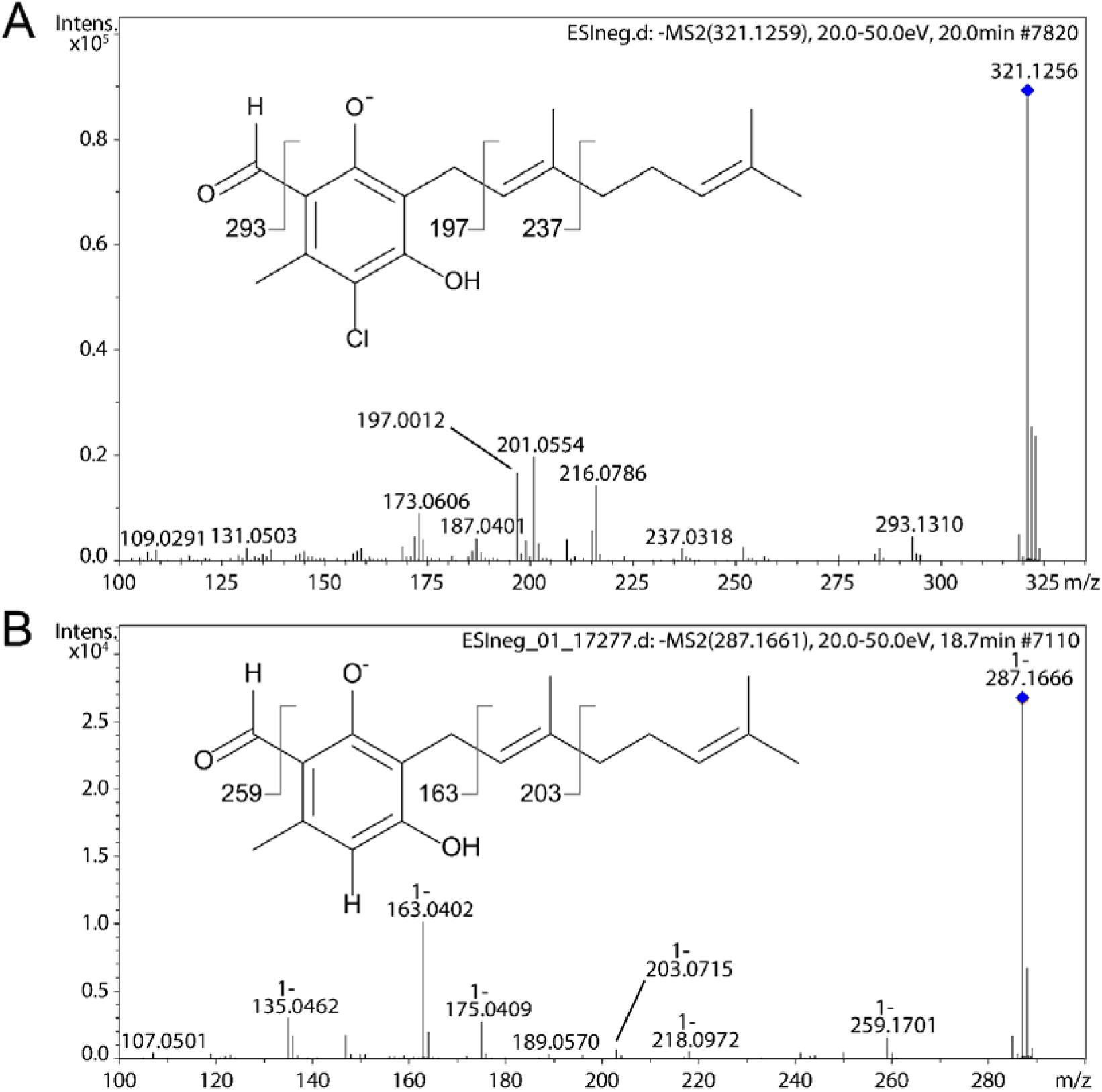
LC-MS/MS fragmentation pattern of, A, a Colletochlorin B standard and, B, a molecule annotated as Colletorin B from the BJNBC-015 strain. Detailed fragmentation patterns are presented in Supplementary File 6.

## 4. Discussion

Fungi are a huge and underestimated reservoir of bioactive natural products. While genetic and chemical manipulations of fungi are common strategies to activate biosynthetic pathways in laboratory conditions, they require extensive trial-and-error. In order to facilitate the discovery of new fungal specialized metabolites, we developed the system described in this study to provide an easily-applicable tool for the heterologous expression of entire secondary metabolite gene clusters in engineered S. cerevisiae. The expression system should facilitate the discovery of new fungal specialized metabolites, especially those produced by silent BGCs, or BGCs that are only expressed at low levels in vitro or uniquely during interactions with the host plant. Apart from the possible discovery of high-value natural products including new medicines or biopesticides, this will provide a better understanding of the ecological role of these molecules, including their contribution to the pathogenesis of plant pathogens. The host organism, Saccharomyces cerevisiae, is a GRAS (generally recognized as safe) organism and is easily cultured on a large scale, while the polycistronic plasmid allows for simultaneous enzyme production and avoids multiple cloning and transformation steps with many vectors for introducing each gene with its own promoter in the engineered recipient strain. In addition, the fluorescent reporter protein VENUS provides a simple way to check the transcription and translation of the polycistronic gene. The production of toxic metabolites may also be possible given the use of the inducible promoters pADH2 and pPCK1, which are activated after the diauxic shift (Harvey et al., 2018), allowing yeast biomass to increase before potentially toxic metabolites start to be produced.

The expression system was firstly validated in yeast by introducing the mScarlet reporter gene into the adapted polycistronic plasmid. We then introduced the genes coding the key and tailoring enzymes (CTC1 to CTC4) of C. higginsianum BGC16 and showed that the resulting yeast cultures heterologously produced Orsellinic acid, Colletorin D acid, Colletochlorin B and D and Colletorin B. The chemical structure of these metabolites validates the correct production and enzymatic activity of the PKS, prenyltransferase, NRPS-like and halogenase enzymes in the heterologous system. Moreover, it demonstrates that ctc1 to ctc4 encode all the enzymes necessary for the biosynthesis of Colletochlorins B and D, Colletorin B and D, Colletorin D acid and Orsellinic acid.

In the first experiment, one bottleneck was the proper functioning of the prenyltransferase CTC4. This enzyme was a predicted UbiA-like membrane-bound prenyltransferase, possessing the typical NDXXDXXXD motif and potentially seven transmembrane domains according to TMHMM (Chang et al., 2021; Krogh et al., 2001). The main hypothesis explaining a non-functional prenyltransferase is the absence of insertion of the protein into the membrane or misfolding followed by rapid degradation of the protein. Although not detected by the signal peptide predictor SignalP (Petersen et al., 2011), this enzyme is predicted to have a plasma-membrane addressing signal by WoLF PSORT (Horton, 2007). Generally, proteins destined to be transported into the membrane are synthesized with a targeting-sequence, usually at the N-terminus and possibly at the C-terminus (Schatz and Dobberstein, 1996). The porcine tescho virus P2A self-cleaving peptide (P2A) has the peculiarity to add 21 amino acids at the C-terminus of the upstream protein and a proline at the N-terminus of the downstream protein (Kim et al., 2011). The presence of these extra residues likely caused the loss of function observed for the prenyltransferase CTC4. An example of non-recognition of an N-terminal signal peptide due to the presence of the P2A-proline was reported previously for an N-myristoylated protein (Hadpech et al., 2018).

Another hypothesis could be that the prenyltransferase is correctly translocated into the membrane but the catalytic site of the enzyme is non-active due to presence of the P2A tail, as was reported by Mattern et al. (2017) for an O-acetyltransferase. However, this explanation is less probable because in that case the prenyltransferase would have been detected by western blot. Whatever the explanation, proper functioning of the prenyltransferase appears to have been prevented by the P2A peptide, because Colletochlorins were successfully produced when the prenyltransferase was expressed from a separate plasmid with a conventional construction involving a strong promoter and terminator, in addition to the polycistronic plasmid containing the other ctc genes. In future experiments, we recommend to check for the presence of predicted signal peptide and transmembrane domains in the enzymes of each pathway of interest and to clone genes encoding this type of enzyme in a separate plasmid. Alternatively, the TEV protease enzyme may be used to cut the P2A C-terminal tail to avoid interference with the enzyme activity.

The use of a polycistronic 2A sequence for the production of heterologous specialized metabolites in yeast was previously used by Beekwilder et al. (2014). They successfully produced β-carotene with a polycistronic plasmid containing three genes separated by the Thosea asigna virus 2A sequence (T2A). Jiao et al. (2018) tried to improve the Beekwilder et al. experimental procedure by studying the order of the genes introduced in the polycistronic plasmid. They concluded that the first gene was more highly expressed than the two following ones. It may thus be better to put the gene encoding the most rate-limiting enzyme at the beginning of the polycistron. Liu et al. (2017) also found that gene expression level progressively decreases with distance from the N-terminus of the polycistron. One limitation of the polycistronic plasmid may therefore be the number of genes introduced. To overcome this issue, we designed two polycistronic plasmids pHYX137 and pHYX138 with different markers of prototrophy, namely LEU2 and URA3, respectively. Distributing the genes of the BGC between these two polycistronic plasmids may allow a more homogeneous expression level for BGCs containing numerous genes.

The successful heterologous production of Colletochlorins demonstrated that four genes of the BGC16 of C. higginsianum are sufficient for the production of these molecules. Eight other genes in the BGC16 were previously assigned to Higginsianins biosynthesis (Tsukada et al., 2020), while the gene CH63R_05474 is a methyltransferase relict that underwent pseudogenization, CH63R_05472 is a putative transcription factor and CH63R_05481 has no characterized function. Two biosynthetic pathways for the production of Colletochlorins were proposed (Figure 6). Various lines of evidence suggest that the second pathway is the most probable. Li et al. (2016) described the cluster involved in LL-Z1272β (Ilicicolin B) synthesis, which contains three genes coding for a PKS StbA, a prenyltransferase StbC and an NRPS-like StbB. These authors showed that the PKS is involved in the formation of Orsellinic acid, which is then converted into Grifolic acid by the prenyltransferase. The NRPS-like StbB is only able to convert the prenylated form of Orsellinic acid (i.e. Grifolic acid) into LL-Z1272β and does not accept Orsellinic acid as a substrate. In the literature, NRPS-like enzymes that require prenylated substrates have been rarely described. Comparison of the Adenylation domains of the NRPS-like accepting Orsellinic acid (ATEG_03630) or only prenylated-orsellinic acid (StbB) as substrate showed differences in their protein sequence. At position 334, ATEG_03630 possesses a leucine and StbB a glycine, while at position 358, essential for ATEG_03630 substrate specificity (Wang and Zhao, 2014), ATEG_03630 has a histidine and StbB a phenylalanine. The C. higginsianum NRPS-like enzyme CTC3 has the same amino acids involved in substrate specificity as StbB, suggesting that it may have a similar substrate specificity towards prenylated Orsellinic acid. Finally, the proposed ability of prenyltransferase CTC4 to accept both DMAPP and GPP moieties as substrates is known to occur in other aromatic prenyltransferases (Chen et al., 2017; Cheng and Li, 2014; Kalén et al., 1990; Suzuki et al., 1994; Swiezewska et al., 1993). Further experiments are now needed to confirm the Colletochlorin biosynthetic pathway, notably by purifying the prenyltransferase and NRPS-like enzymes for assessing their substrate specificity.

The Colletochlorins were previously isolated from a C. higginsianum mutant with a partially deficient COMPASS complex (Dallery et al., 2019) and several of them were shown previously to be biologically active. For example, Colletorin B and Colletochlorin B displayed moderate herbicidal, antifungal and antibacterial activities towards Chlorella fusca, Ustilago violacea, Fusarium oxysporum, and Bacillus megaterium (Hussain et al., 2015), while Colletochlorin B had a significant antibacterial effect against Bacillus subtilis (minimum inhibitory concentration, 2 μg·mL^-1^) (Kemkuignou et al., 2022).

## 5. Conclusions

Our findings demonstrate the utility of this synthetic biology tool for the metabolic engineering of yeast to produce fungal metabolites from BGCs of interest in bulk liquid cultures. This is a prerequisite for subsequent structural characterization and bioactivity profiling of SM products from BGCs that are otherwise silent in their native organisms when cultured in laboratory conditions.

## Acknowledgments

This work has benefited from the support of IJPB’s “Plant Observatory – Chemistry and Metabolism” platform. The authors would like to thank Axel A. Brakhage and Maria Stroe (Hans Knoll Institute, Jena, Germany), Nancy DaSilva (UC Irvine, California) and Verena Siewers (Chalmers Univ. of Technology, Gothenburg, Sweden) for kindly providing the pV2A-T plasmid, the BJ5464-NpgA strain and the XI-3-pCfb2904-bccpr1 plasmid, respectively. The EasyClone-MarkerFree Vector Set was a gift from Irina Borodina (Addgene kit #1000000098).

## 7. Contribution statement

Conceptualization: JFD, RJO, MV; Methodology: JFD, AGK; Resources: JO, GM; Investigation: AGK, JFD, JCT, GLG, JV, KS; Formal analysis: AGK, JCT, GLG, JFD; Visualization: AGK, JFD, JCT, JV, KS; Writing – Original Draft: AGK, JFD, RJO, MV, JCT; Writing – Review & Editing: JFD, RJO, MV, AGK; Supervision: JFD, RJO, MV, JO, GM; Funding acquisition: JFD, MV, GM, JO; Project administration: JFD.

## 9. Supplementary Files

Supplementary File 1: List of strains used in this study.

Supplementary File 2: Maps of the plasmids pHYX137 and pHYX138.

Supplementary File 3: List of the primers used in this study.

Supplementary File 4: Sequences of the CTC genes codon-adapted for Saccharomyces cerevisiae.

Supplementary File 5: List of the plasmids used in this study.

Supplementary File 6: Fragmentation patterns of Colletochlorin B standard and Colletorin B.

## Abbreviations

BGC: biosynthetic gene cluster
CDS: coding sequence
DAPI: 4’,6-diamidino-2-phenylindole
DMAPP: dimethylallyl pyrophosphate
ELSD: evaporative light scattering detector
GPP: geranyl pyrophosphate
HPLC: high-performance liquid chromatography
IVA: in vivo assembly
LC: liquid chromatography
MS: mass spectrometry
NLS: nuclear localization signal
NRPS: non-ribosomal peptide synthetase
OA: Orsellinic acid
OSMAC: one strain many compounds
PCR: polymerase chain reaction
PDA: potato dextrose agar
PKS: polyketide synthase
SM: secondary metabolite
TAR: transformation-assisted recombination
TEV: Tobacco etch virus
UV: ultraviolet
YNB: yeast nitrogen broth
YPD: yeast extract peptone dextrose.

