## Supplementary File 6 for "Yeast-based heterologous production of the Colletochlorin family of fungal secondary metabolites"

### Slide 1
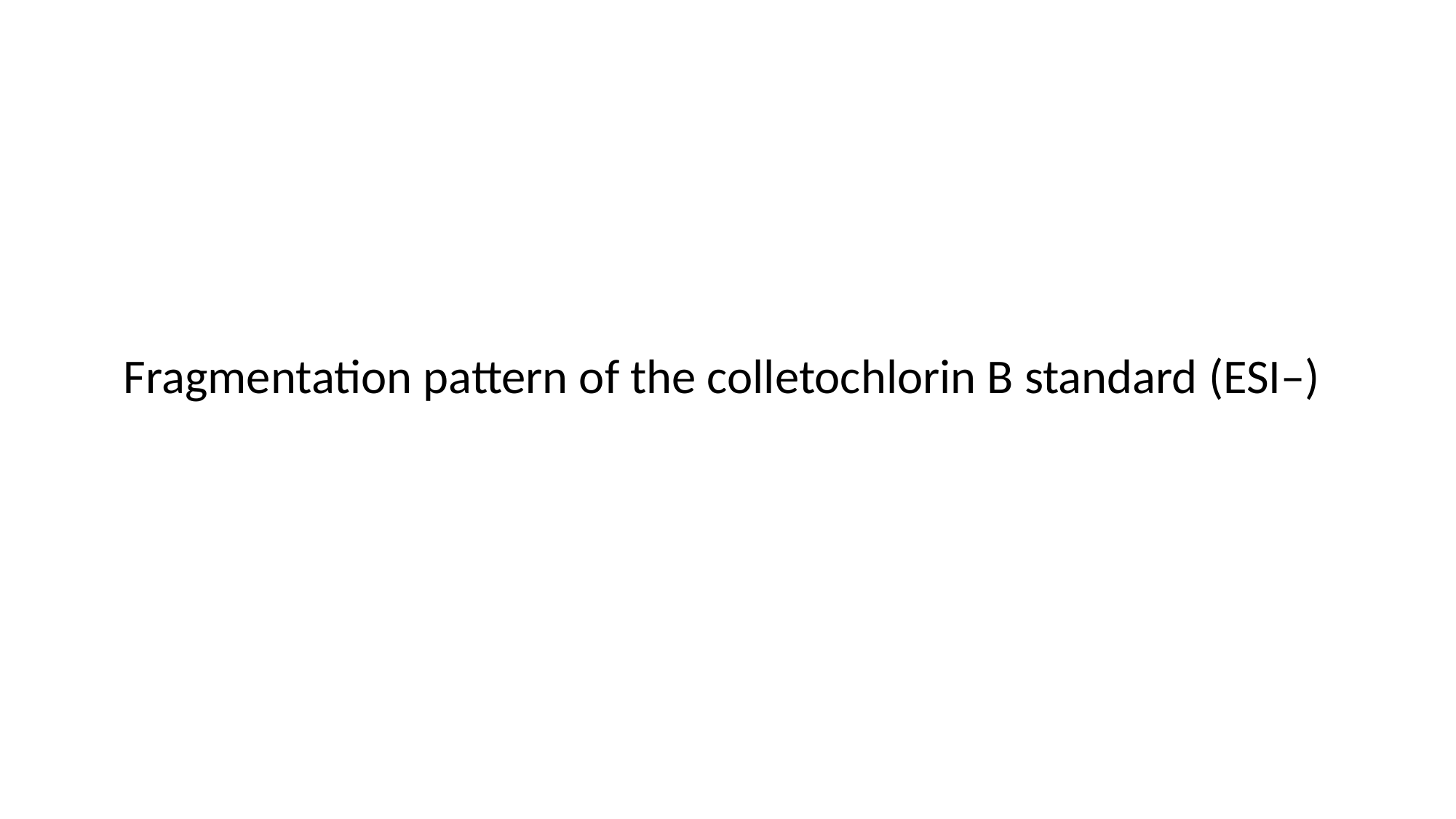

Fragmentation pattern of the colletochlorin B standard (ESI–)

### Slide 2
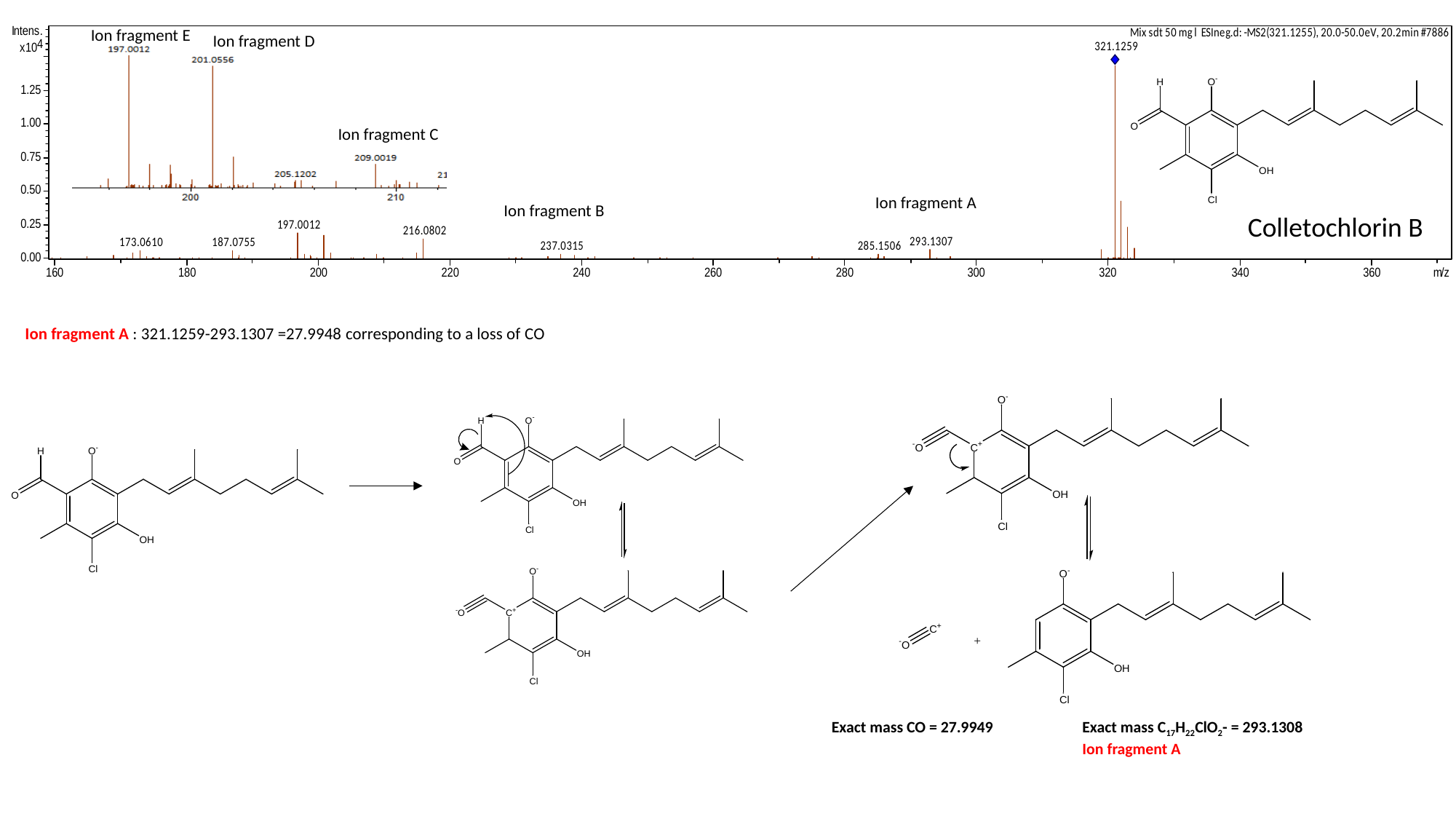

Ion fragment E
Ion fragment D
Ion fragment C
Ion fragment A
Ion fragment B
Colletochlorin B
Ion fragment A : 321.1259-293.1307 =27.9948 corresponding to a loss of CO
Exact mass C_17_H_22_ClO_2_- = 293.1308
Ion fragment A
Exact mass CO = 27.9949

### Slide 3
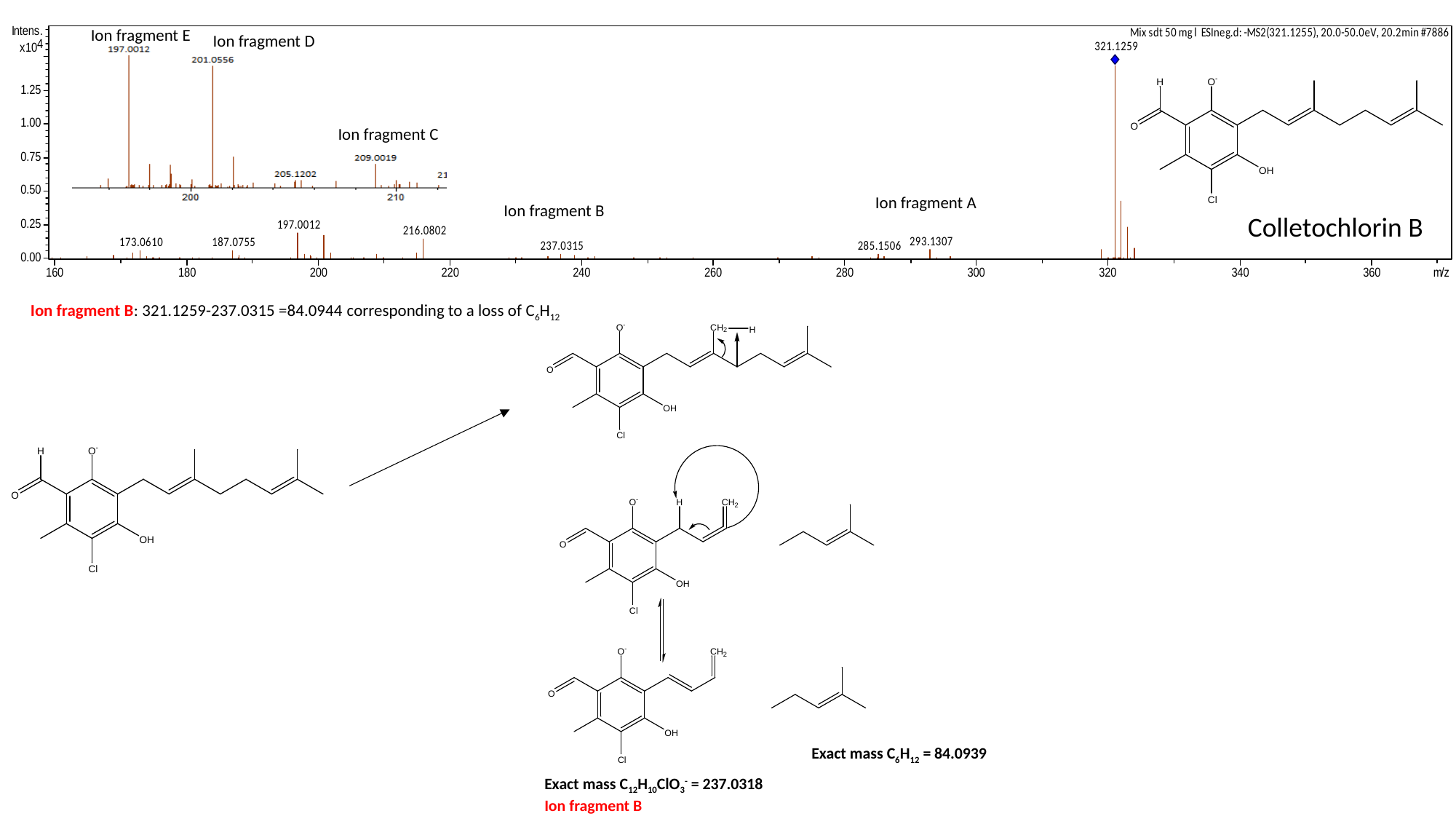

Ion fragment E
Ion fragment D
Ion fragment C
Ion fragment A
Ion fragment B
Colletochlorin B
Ion fragment B: 321.1259-237.0315 =84.0944 corresponding to a loss of C_6_H_12_
Exact mass C_6_H_12_ = 84.0939
Exact mass C_12_H_10_ClO_3_^-^ = 237.0318
Ion fragment B

### Slide 4
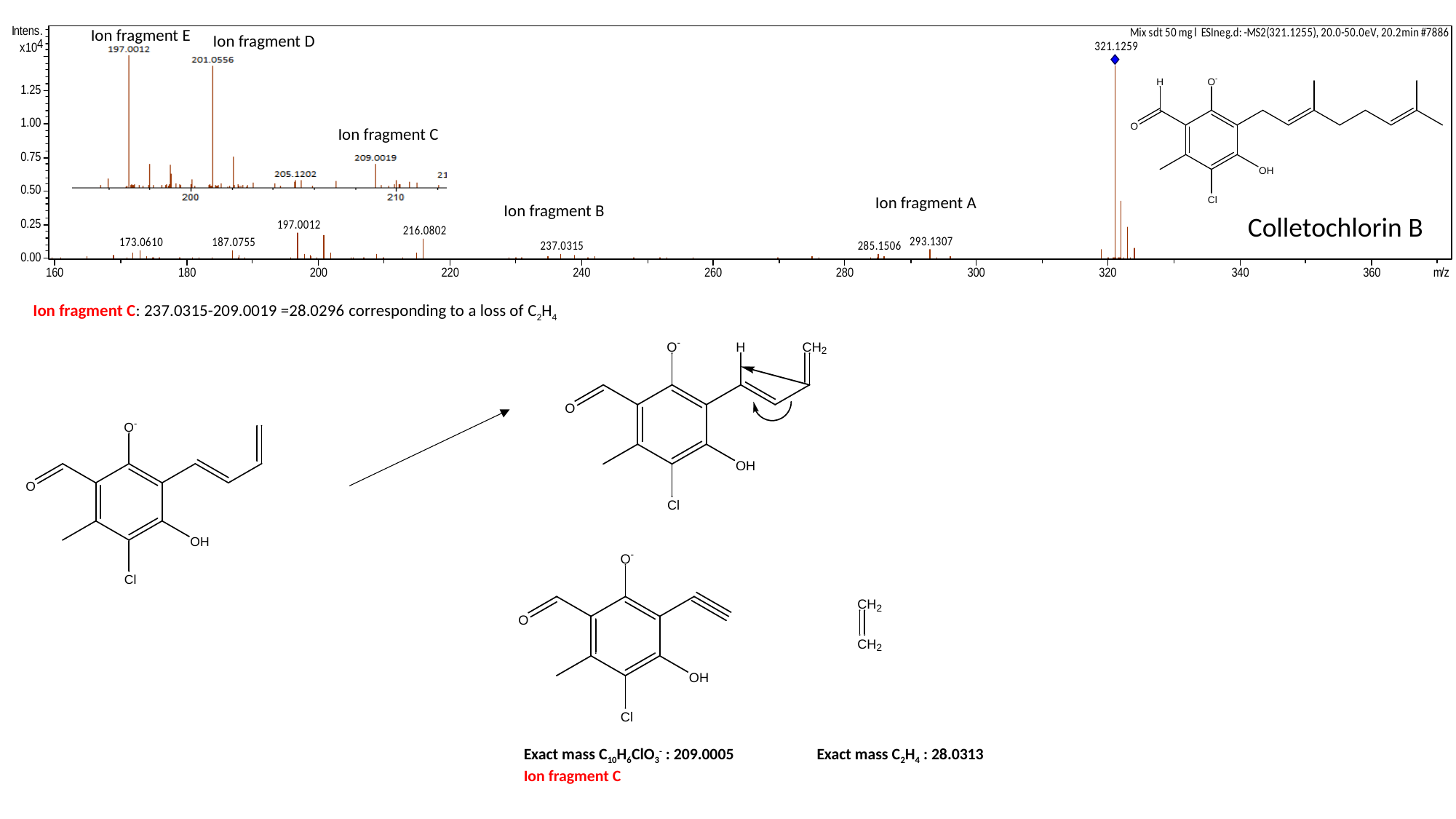

Ion fragment E
Ion fragment D
Ion fragment C
Ion fragment A
Ion fragment B
Colletochlorin B
Ion fragment C: 237.0315-209.0019 =28.0296 corresponding to a loss of C_2_H_4_
Exact mass C_10_H_6_ClO_3_^-^ : 209.0005
Ion fragment C
Exact mass C_2_H_4_ : 28.0313

### Slide 5
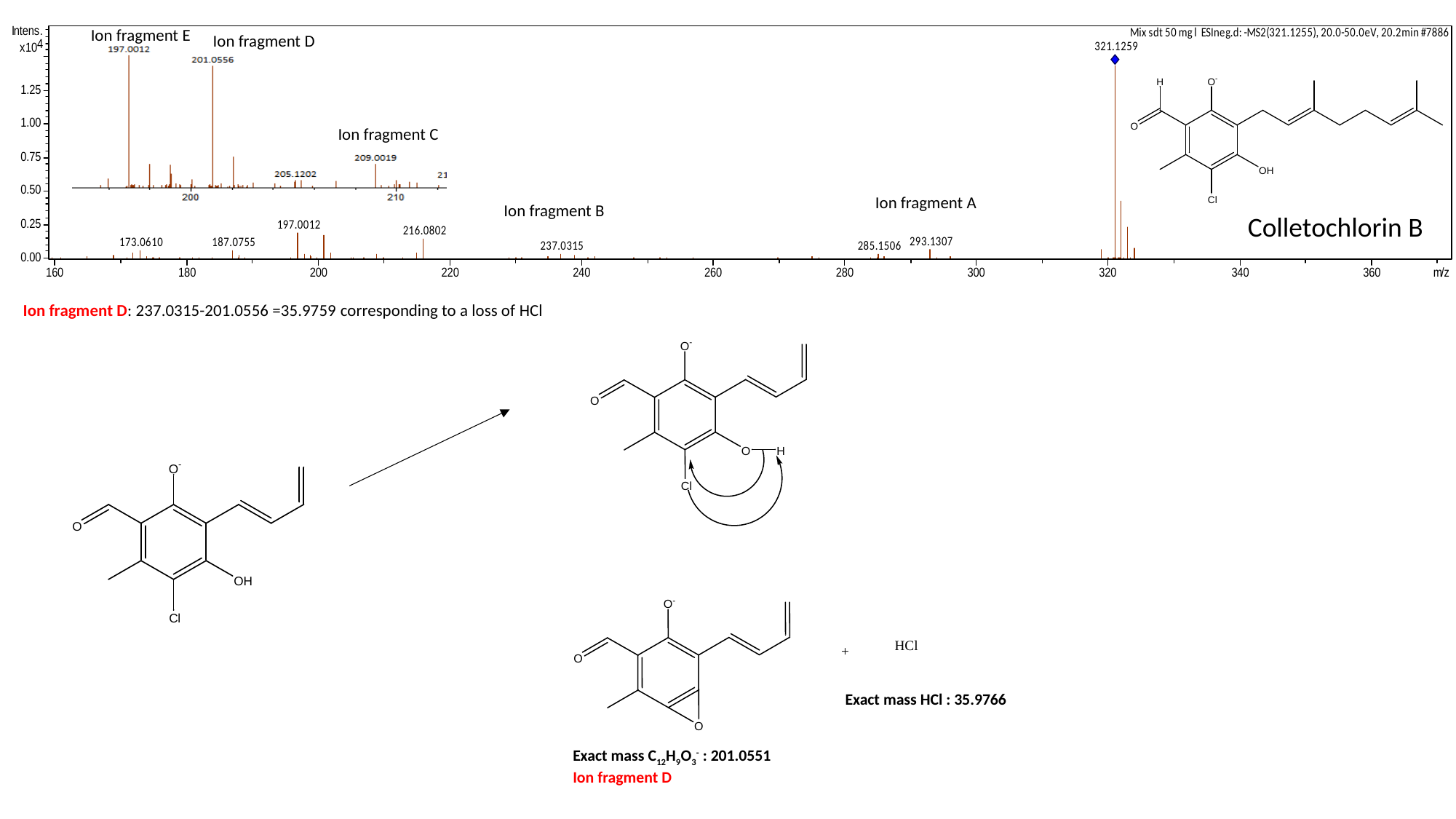

Ion fragment E
Ion fragment D
Ion fragment C
Ion fragment A
Ion fragment B
Colletochlorin B
Ion fragment D: 237.0315-201.0556 =35.9759 corresponding to a loss of HCl
Exact mass HCl : 35.9766
Exact mass C_12_H_9_O_3_^-^ : 201.0551
Ion fragment D

### Slide 6
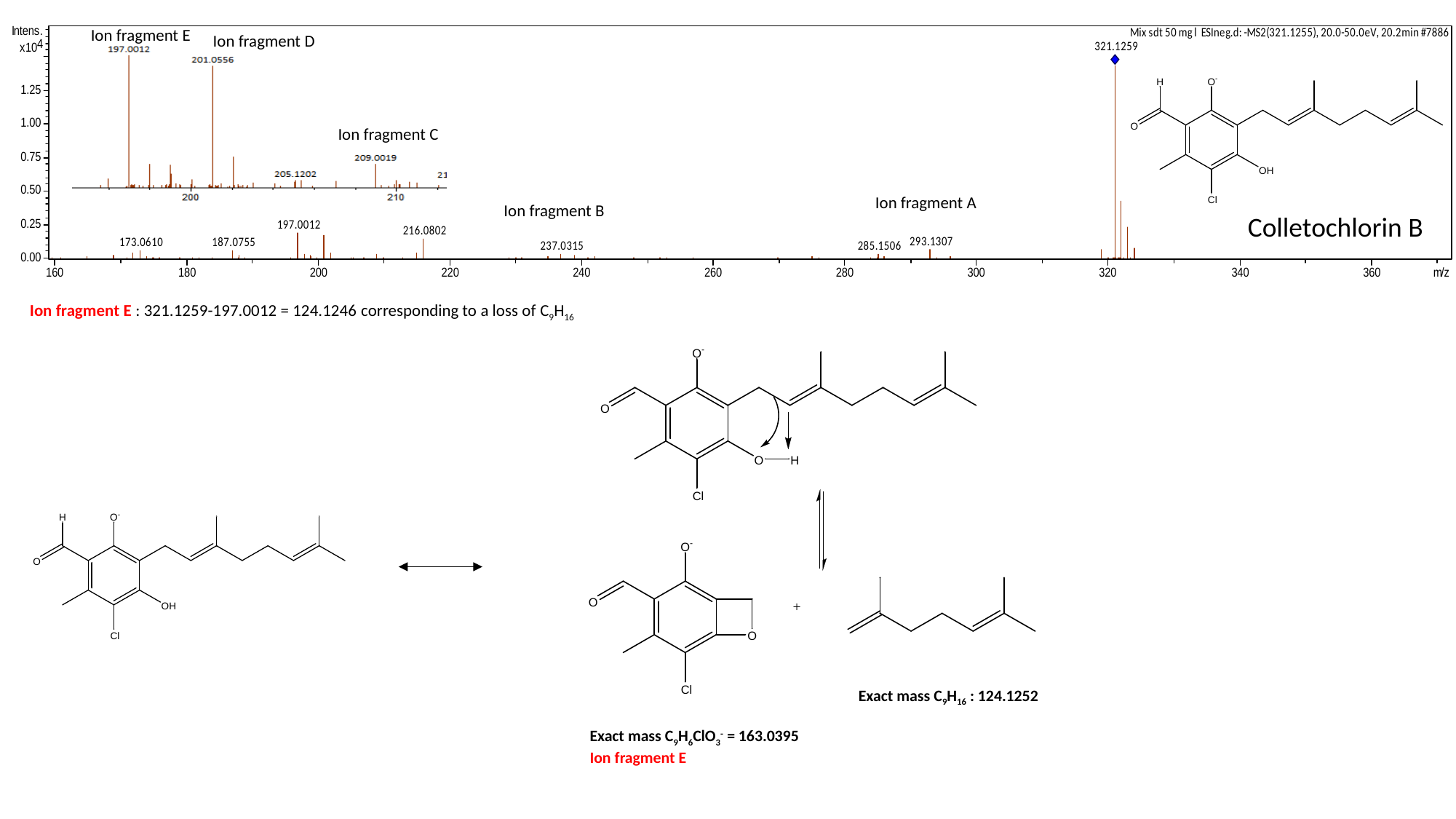

Ion fragment E
Ion fragment D
Ion fragment C
Ion fragment A
Ion fragment B
Colletochlorin B
Ion fragment E : 321.1259-197.0012 = 124.1246 corresponding to a loss of C_9_H_16_
Exact mass C_9_H_16_ : 124.1252
Exact mass C_9_H_6_ClO_3_^-^ = 163.0395
Ion fragment E

### Slide 7
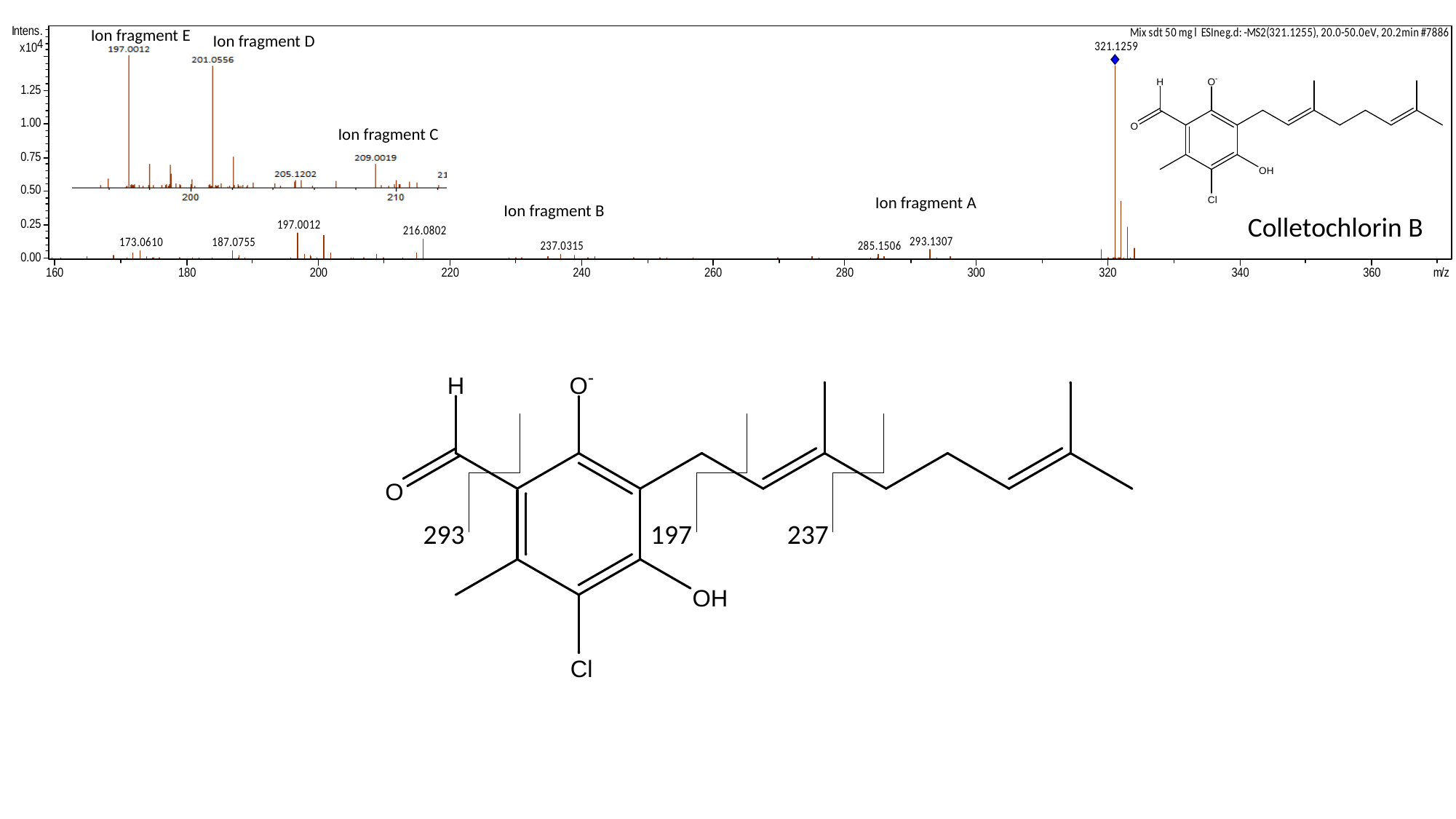

Ion fragment E
Ion fragment D
Ion fragment C
Ion fragment A
Ion fragment B
Colletochlorin B
293
197
237

### Slide 8
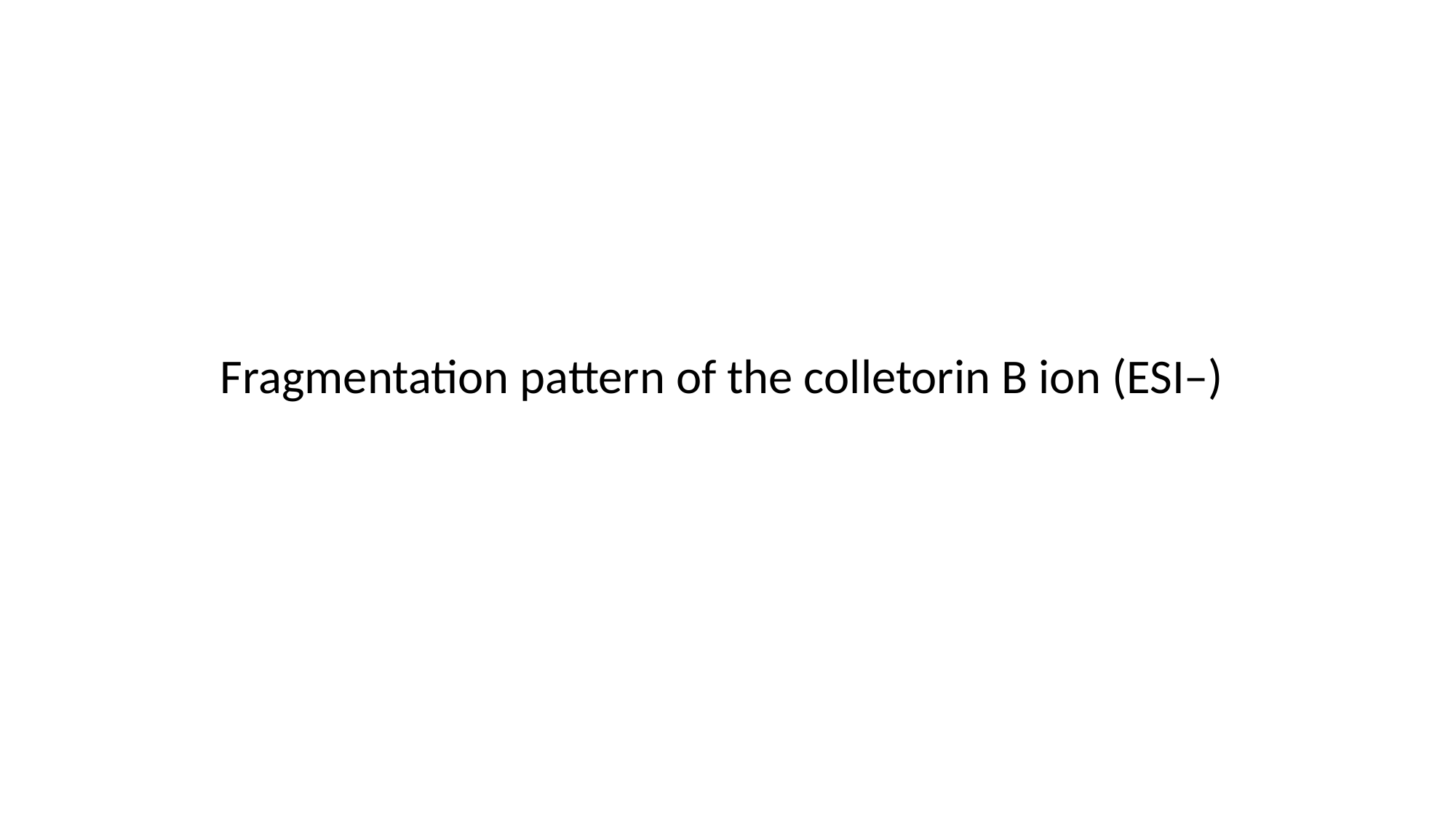

Fragmentation pattern of the colletorin B ion (ESI–)

### Slide 9
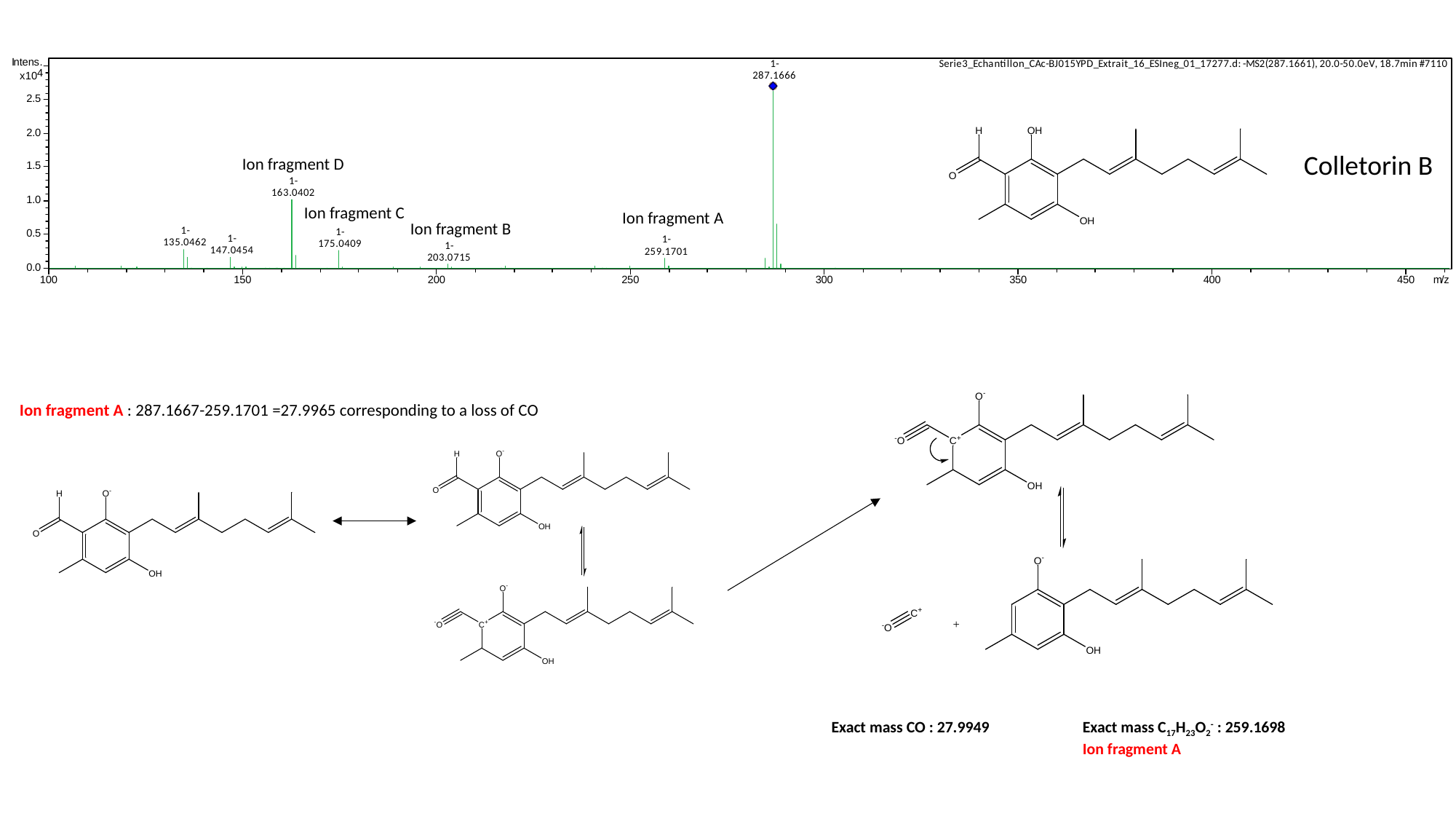

Colletorin B
Ion fragment D
Ion fragment C
Ion fragment A
Ion fragment B
Ion fragment A : 287.1667-259.1701 =27.9965 corresponding to a loss of CO
Exact mass C_17_H_23_O_2_^-^ : 259.1698
Ion fragment A
Exact mass CO : 27.9949

### Slide 10
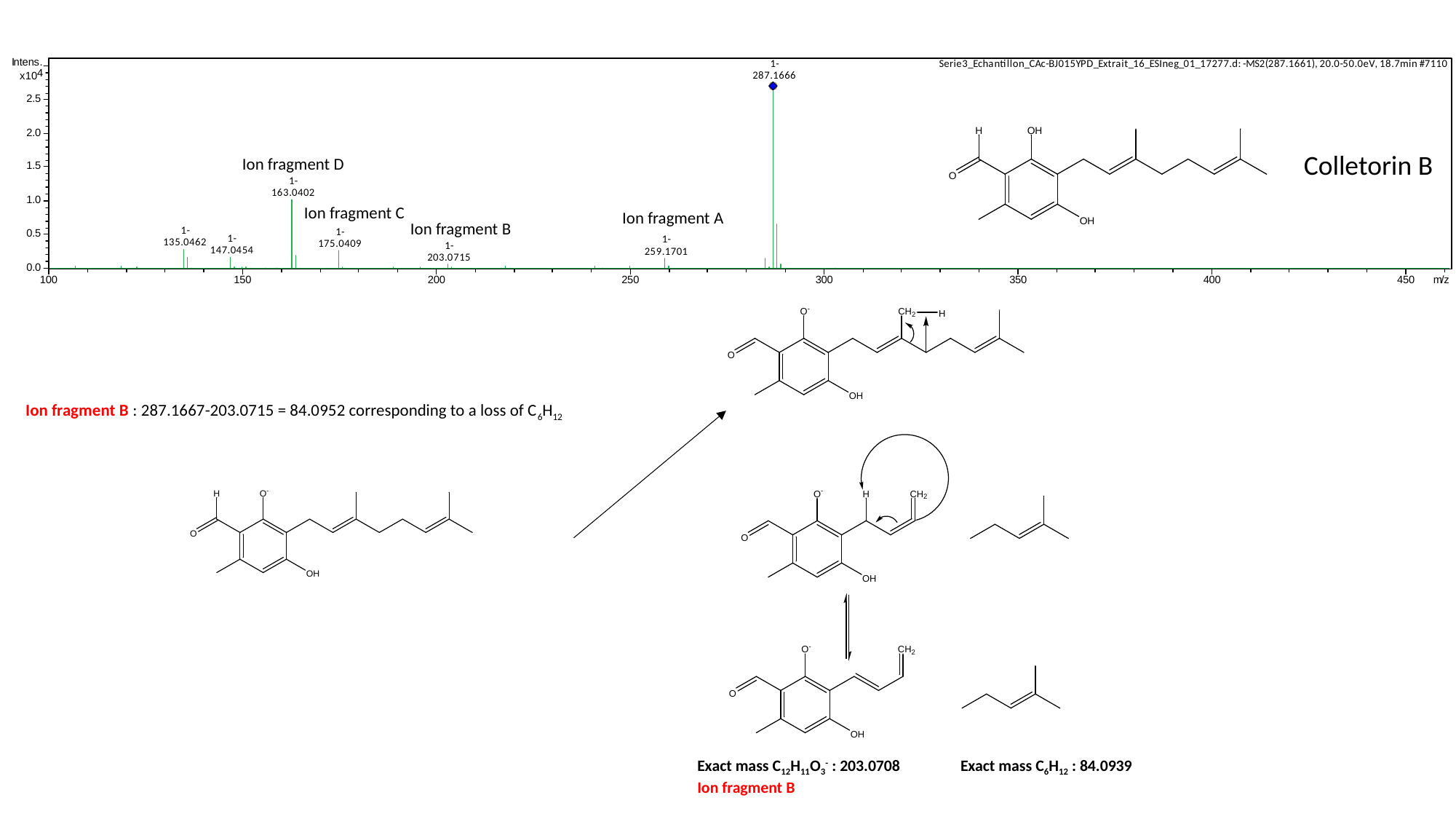

Colletorin B
Ion fragment D
Ion fragment C
Ion fragment A
Ion fragment B
Ion fragment B : 287.1667-203.0715 = 84.0952 corresponding to a loss of C_6_H_12_
Exact mass C_12_H_11_O_3_^-^ : 203.0708
Ion fragment B
Exact mass C_6_H_12_ : 84.0939

### Slide 11
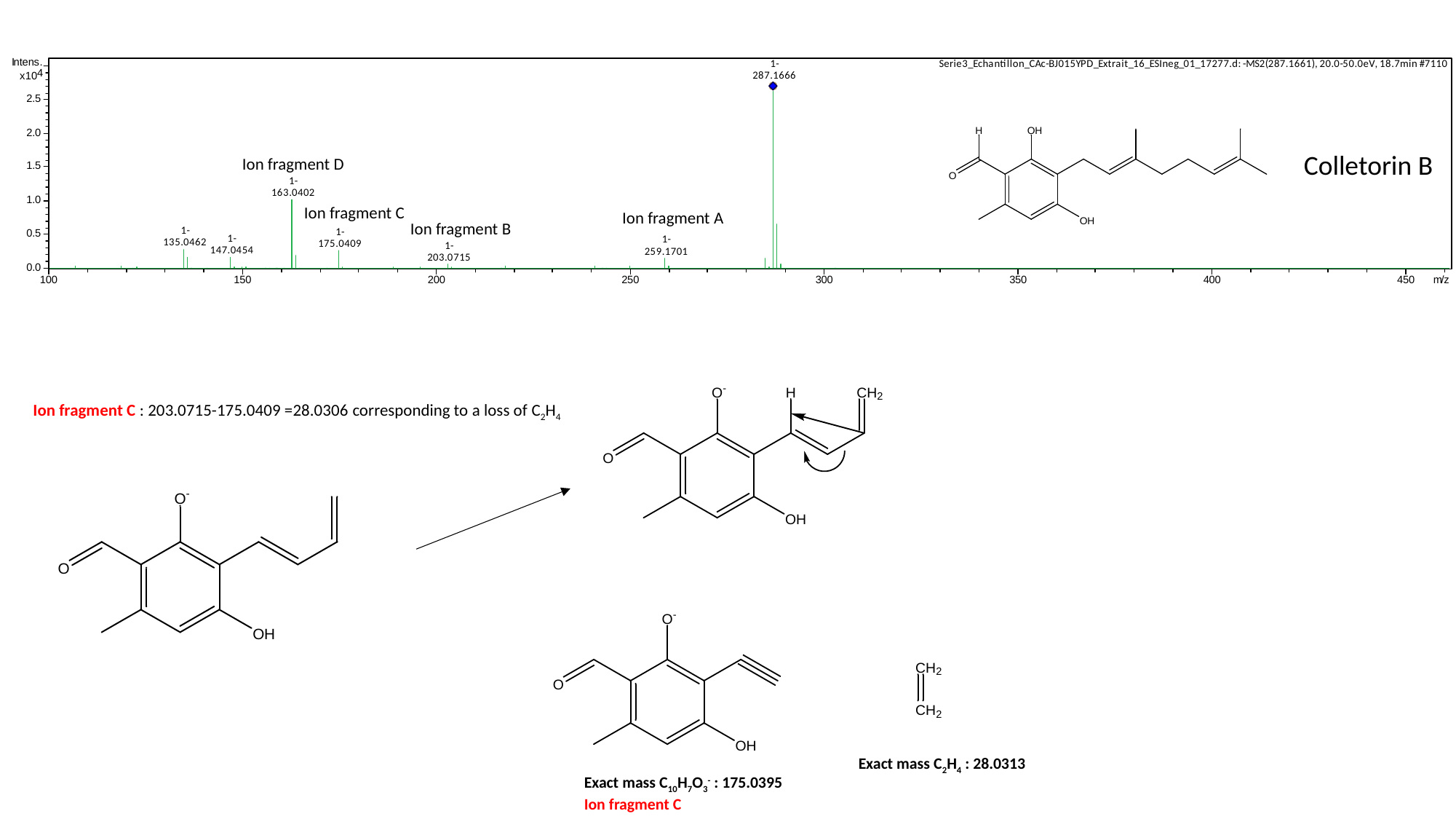

Colletorin B
Ion fragment D
Ion fragment C
Ion fragment A
Ion fragment B
Ion fragment C : 203.0715-175.0409 =28.0306 corresponding to a loss of C_2_H_4_
Exact mass C_2_H_4_ : 28.0313
Exact mass C_10_H_7_O_3_^-^ : 175.0395
Ion fragment C

### Slide 12
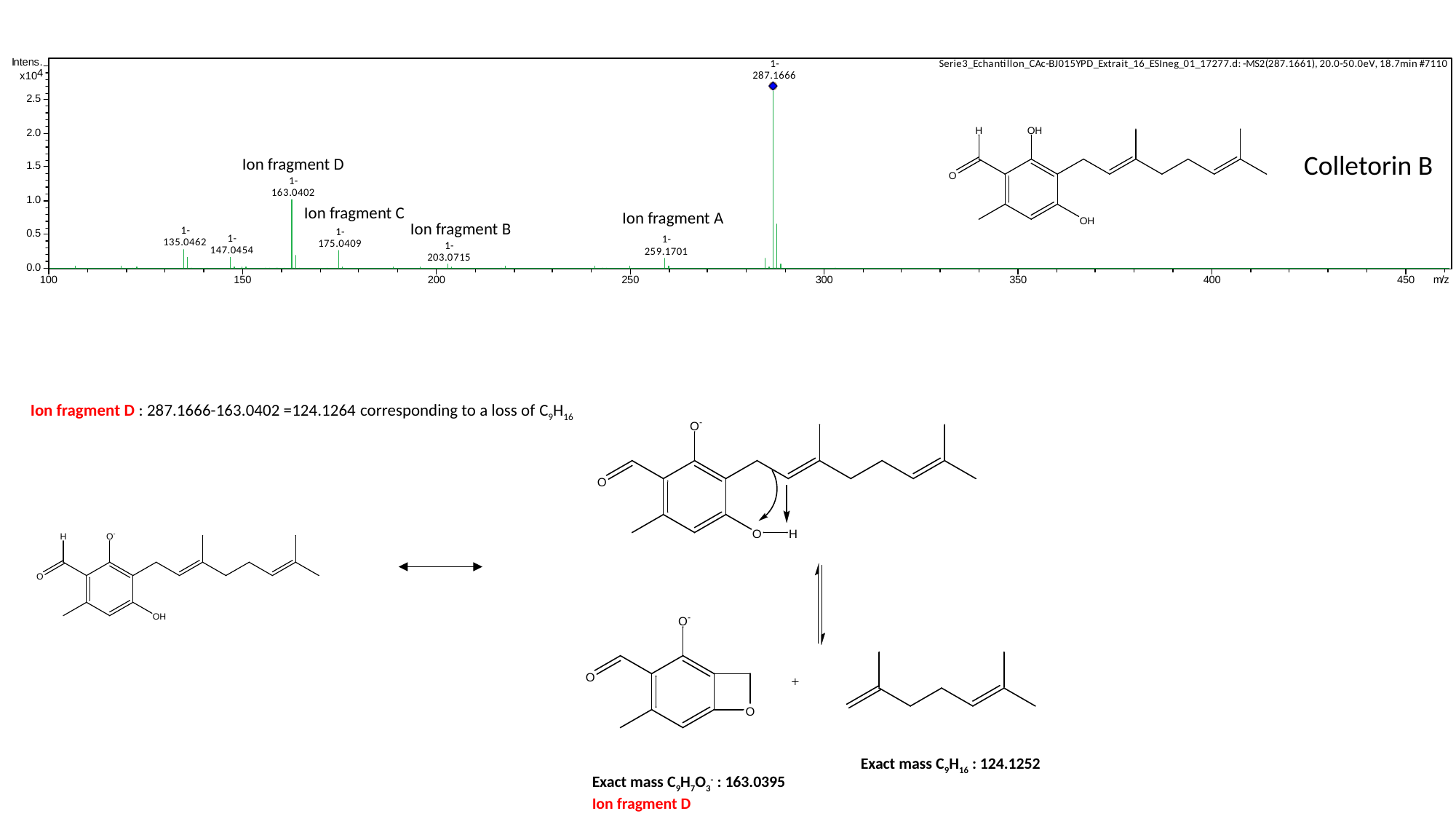

Colletorin B
Ion fragment D
Ion fragment C
Ion fragment A
Ion fragment B
Ion fragment D : 287.1666-163.0402 =124.1264 corresponding to a loss of C_9_H_16_
Exact mass C_9_H_16_ : 124.1252
Exact mass C_9_H_7_O_3_^-^ : 163.0395
Ion fragment D

### Slide 13
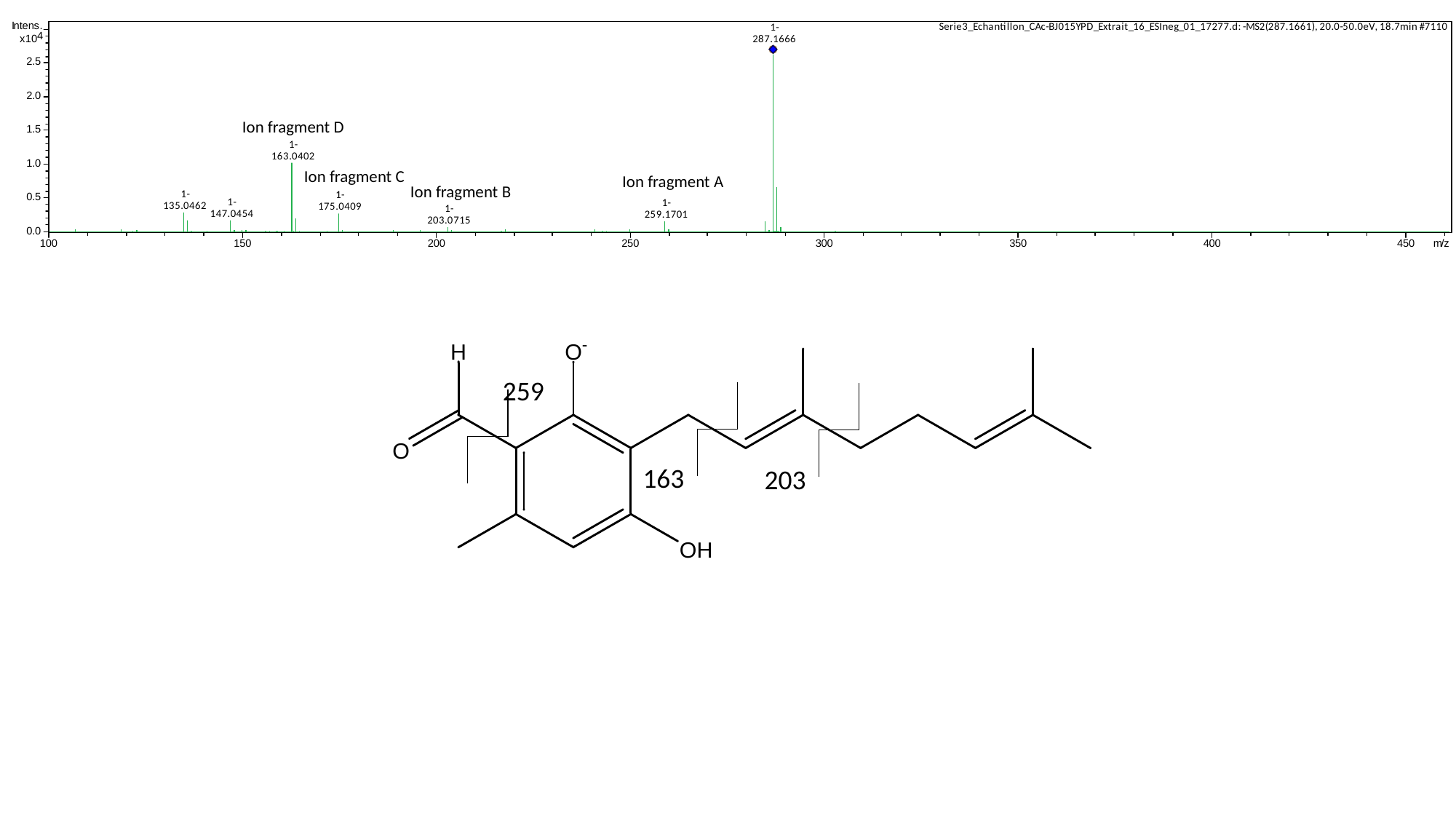

Ion fragment D
Ion fragment C
Ion fragment A
Ion fragment B
259
163
203
