## Supplementary figures and images for "Yeast-based heterologous production of the Colletochlorin family of fungal secondary metabolites"

### Supplementary File 2

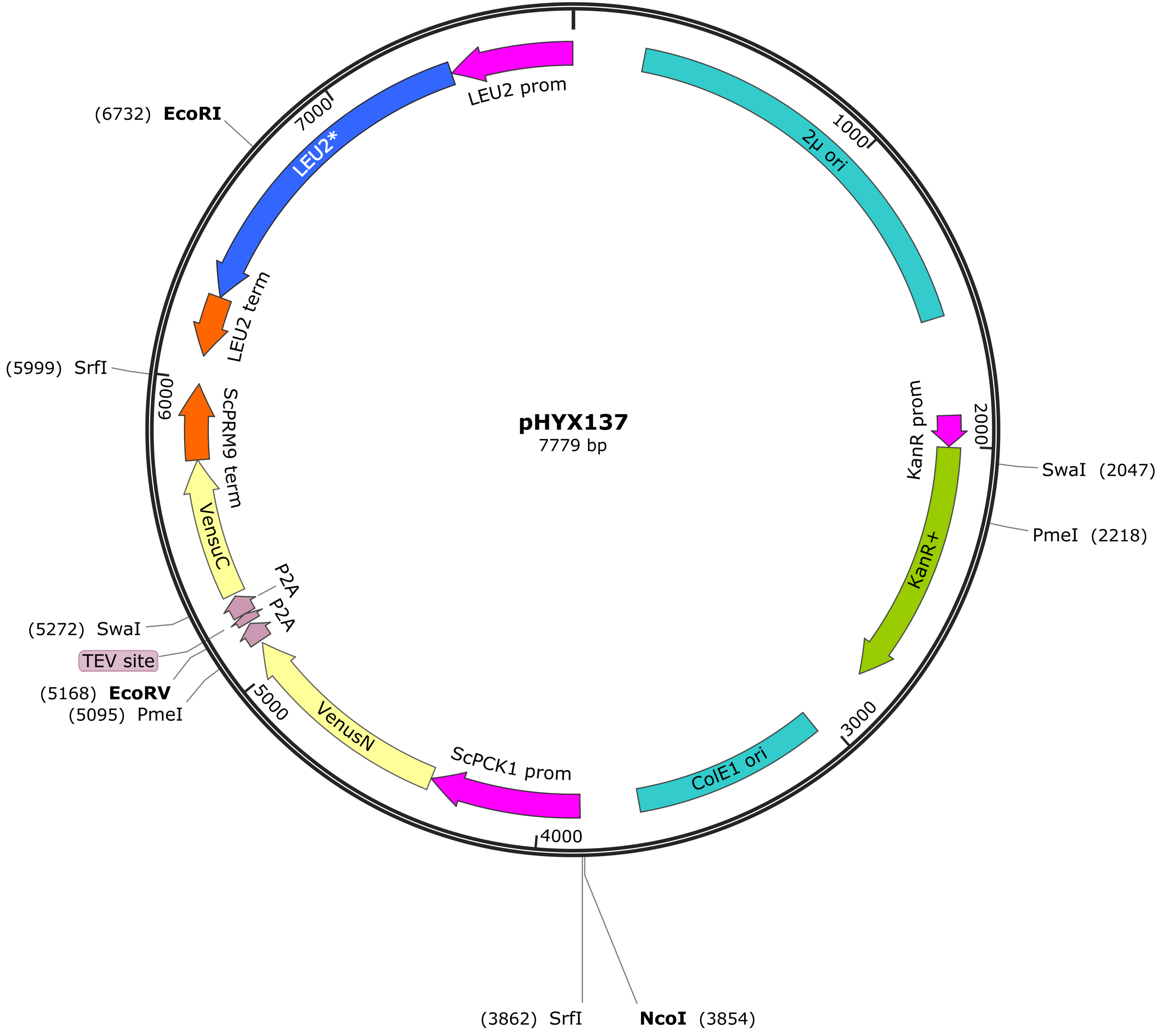

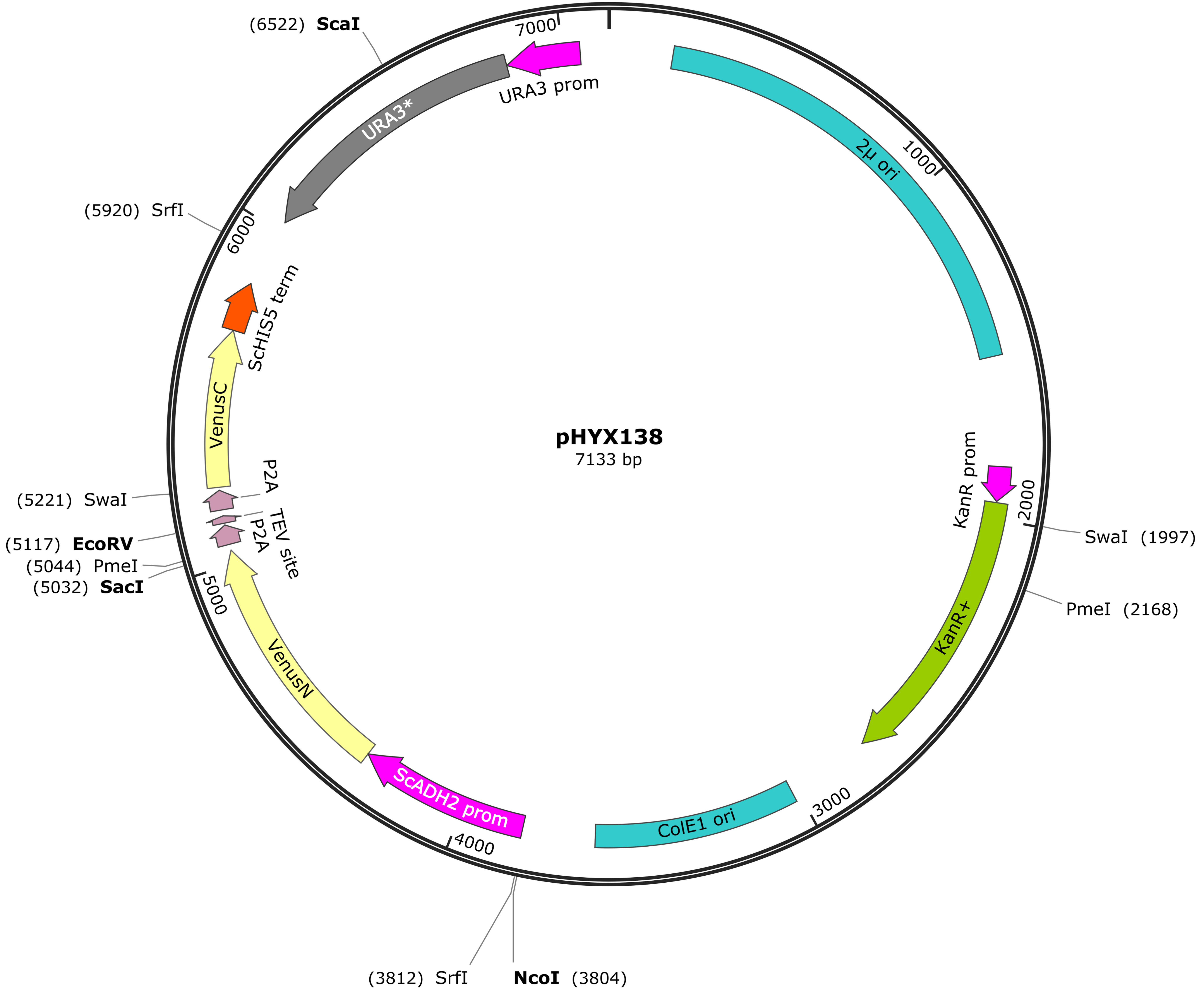
